# Identification of a Broadly Fibrogenic Macrophage Subset Induced by Type 3 Inflammation in Human and Murine Liver and Lung Fibrosis

**DOI:** 10.1101/2022.07.01.498017

**Authors:** Thomas Fabre, Alexander M. S. Barron, Stephen M. Christensen, Shoh Asano, Marc H. Wadsworth, Xiao Chen, Ju Wang, James McMahon, Frank Schlerman, Alexis White, Kellie Kravarik, Andrew J. Fisher, Lee A. Borthwick, Kevin M. Hart, Neil C. Henderson, Thomas A. Wynn, Ken Dower

## Abstract

Macrophages are central orchestrators of the tissue response to injury, with distinct macrophage activation states playing key roles in the progression and resolution of fibrosis. Identifying the unique fibrogenic macrophages that are found in human fibrotic tissues could lead to new and more effective treatments for fibrosis. Here we used human liver and lung single cell RNA sequencing datasets to identify a unique subset of CD9^+^ TREM2^+^ macrophages expressing SPP1, GPNMB, FABP5, and CD63 with strong pro-fibrotic activity. This population was validated across orthogonal techniques, species and tissues. These macrophages were enriched at the outside edges of scarring adjacent to activated mesenchymal cells, and in the fibrotic niche across species and organs. Neutrophils producing the type 3 cytokines GM-CSF and IL-17A, and expressing MMP9, which participates in the activation of TGF-β1, clustered with these scar-associated macrophages. Using *in vitro* primary human cell assays, we determined that GM-CSF, IL-17A and TGF-β1 drive the differentiation of these scar-associated macrophages, and that co-culture of monocyte-derived macrophages with hepatic stellate cells and TGF-β1 augmented type 1 collagen deposition. *In vivo* blockade of GM-CSF, IL-17A or TGF-β1 with small or large molecules reduced scar-associated macrophage expansion and fibrosis in multiple models of hepatic and pulmonary fibrosis. Our work demonstrates that a specific scar-associated macrophage population is linked with fibrosis across species and tissues. It further provides a strategy for unbiased discovery, triage and preclinical validation of therapeutic targets within this fibrogenic macrophage population.

## Introduction

Organ fibrosis represents a major cause of morbidity and mortality worldwide (*1*). Fibrosis can arise from multiple etiologies and affects virtually any organ (*2*). Prominent among fibrotic diseases are non-alcoholic steatohepatitis (NASH), which is the fastest growing cause of liver fibrosis (*3*), and idiopathic pulmonary fibrosis (IPF), which although relatively rare, is the prototypical progressive-fibrosing interstitial lung disease (*4*). In both diseases, inflammation shapes the complex cellular architecture and interactions and phenotypes of scar-associated cells within the fibrotic niche (*1, 3, 5-9*). However, it remains unclear as to how much of this cellular dialogue is organ- or disease-specific (*3, 5, 6*).

Macrophages are a critical effector population contributing to both fibrosis progression and resolution (*7, 10*). Recently, ‘omics data have revealed similar scar-associated macrophages (SAMs) expressing TREM2 and/or CD9 in hepatic and pulmonary fibrosis (*11–21*). CD9^+^TREM2^+^ macrophages are also associated with Alzheimer’s disease and are enriched in obese adipose tissue and atherosclerotic lesions across species, suggesting that these markers may be expressed by several myeloid populations with non-overlapping functions (*21–24*). SAMs likely originate from monocytes, localize to the scar, and induce myofibroblast activation (*7, 10-12, 15, 25, 26*). Multiple factors influence monocyte-to-macrophage differentiation including niche and inflammatory signals (*7, 27*). Fibrosis-associated inflammation can vary depending on the disease and model, but in NASH and IPF is characterized by a mixture of type 1, 2, 3 and innate cytokines (*1, 8, 9*). Despite these advances, SAMs from across organs, fibrotic diseases and species have not been precisely defined or directly compared, and the factors influencing SAM differentiation from monocytes remain unclear.

Here we applied ‘omics data and murine preclinical models of lung and liver fibrosis to identify a “core” fibrotic SAM phenotype characterized by expression of *CD9 and TREM2, but also SPP1*, *GPNMB*, *FABP5* and *CD63*. We find that this level of resolution was needed to accurately identify this population. These “core” SAMs were enriched in both lung and liver fibrosis across species. RNA velocity (pseudotime trajectory) analysis suggested that they originated from infiltrating monocytes and represented an intermediate differentiation state. Tissue mapping revealed that SAMs were enriched at the edges of scarring in association with myofibroblasts and IL-17^+^GM-CSF^+^MMP9^+^ neutrophils. SAMs expressed a variety of pro-fibrogenic mediators including chemokines, tissue remodeling molecules and fibroblast-activating factors such as osteopontin (*SPP1*). *In vivo* and *in vitro* we showed that type 3 cytokines, IL-17A, GM-CSF and TGF-β, drives the maturation of SAMs, and that blocking these factors reduces SAM formation and fibrosis severity. Together, our results identified a core SAM population in lung and liver fibrosis that localized to the fibrotic niche and differentiates from monocytes in response to IL-17A, GM-CSF and TGF-β. Targeting SAMs therefore represents a promising strategy to treat human fibrotic diseases.

## Materials and Methods

### Mice

All experimental procedures were performed at Pfizer (Cambridge, MA) and approved by the Institutional Animal Care and Use Committee. The mice were maintained in a controlled environment with a temp 70-72° F, humidity 30-70%, with a photo cycle of 12 hours of light and 12 hours of dark. They were provided with PicoLab rodent diet 20 (5053, Purina Mills, Inc., Lab Diet, St Louis, MO, unless otherwise indicated for high fat diets) and standard drinking water *ad libitum*. C57 BL/6 male mice (N=8/group) 8-10 weeks of age (The Jackson Laboratory, Bar Harbor, Maine) were dosed 2 mL/Kg intraperitoneally (IP) twice weekly with either olive oil (N=8) or CCl_4_ (CCl_4_:olive oil, 1:3, [v:v], Sigma-Aldrich, St Louis, MO) for 4 weeks. Mice were euthanized with C02 at study end. For bleomycin lung injury, on day 0 animals were intra-tracheally challenged with 75 μg of Bleomycin (EMD Millipore, Burlington, MA) in a volume of 25 μl. 7 days later, mice are therapeutically dosed with vehicle, ALK5 inhibitor (SB-525334) for a duration of 14 days. At study end mice are anesthetized with isoflurane, exsanguinated, and euthanized *via* cervical dislocation. For pre-clinical NASH studies, C57 BL/6 male mice, 6-8 weeks of age (The Jackson Laboratory, Bar Harbor, ME) were fed either control (D09100304) or high fat (D09100310) diet (Research Diets, New Brunswick, NJ) for 34 weeks to induce disease phenotype. Shear wave elastography was performed prior to therapeutic treatment and animals were randomized into treatment groups based on average liver stiffness. Following randomization animals were singly housed except for control animals which were group housed. Animals in the control group (N=8) were maintained on control chow through week 44. HFD treatment groups included: IgG1 isotype control (N=13, 10 mg/kg, PF-00713222-0010), TGF-β Ab 1D11 (N=13, 10 mg/kg, BP0057 lot#704719J1, BioXCell, West Lebanon, NH).

### Human lung and liver sections

The study was approved by the NHS national research ethics service under approval numbers 16/NE/0230 and 11/NE/0291. It utilized non-diseased human donor lungs that were not used in clinical transplantation and when consent from the donor next of kin was given for organ research. Explant IPF lung tissue was collected from patients undergoing either double or single lung transplantations at the Freeman Hospital, Newcastle upon Tyne.

Anonymized human NASH liver sections were acquired from BioIVT.

Characteristics of these patients are presented in Supplemental Table 1.

### Single-cell RNA sequencing and analysis

#### Single Cell RNA Sequencing

Cells isolated from murine livers were stained with 30 μg/mL Hoechst 33342 (Miltenyi) and Zombie NIR viability dye (diluted 1:250 in PBS; Biolegend) before FACS sorting (Sony SH800) to obtain live, single cells. These were methanol fixed based on the 10x Genomics recommended fixation protocol and stored at - 80°C before encapsulation and barcoding on a Chromium Controller (10x Genomics, Pleasanton, CA). 3’ sequencing library construction followed the protocols for 10x Genomics Single Cell version 2 or version 3.1. Samples from some experiments were lipid hashed as described by McGinnis *et al.* before encapsulation except using commercially available 5’ or 3’ cholesterol-TEG tagged oligos as the anchor and co-anchor. For these samples, sequencing libraries were constructed as described previously (*28*). Libraries were sequenced on Illumina NextSeq 500/550.

#### Data Preprocessing & Quality Control

Gene expression matrices from Ramachandran et al. (GSE136103, (*11*)), MacParland et al. (HCA project SingleCellLiverLandscape, (*29*)), Reyfman et al. (GSE122960, (*16*)), Morse et al. (GSE128033, (*18*)), Valenzi et al. (GSE128169, (*30*)), Habermann et al. (GSE135893, (*31*)), and Tsukui et al. (GSE132771, (*32*)) were processed using the Seurat (v4.0.3). Ribosomal genes were removed, and cells were selected using cutoffs of minimum 500 genes per cell and maximum 10% mitochondrial reads. Genes per cell, transcript counts per cell and mitochondrial read percentage per cell are presented in Supplemental Figures 1, 2 and 3 (for the human liver, lung and myeloid atlases, respectively).

**Figure 1.**
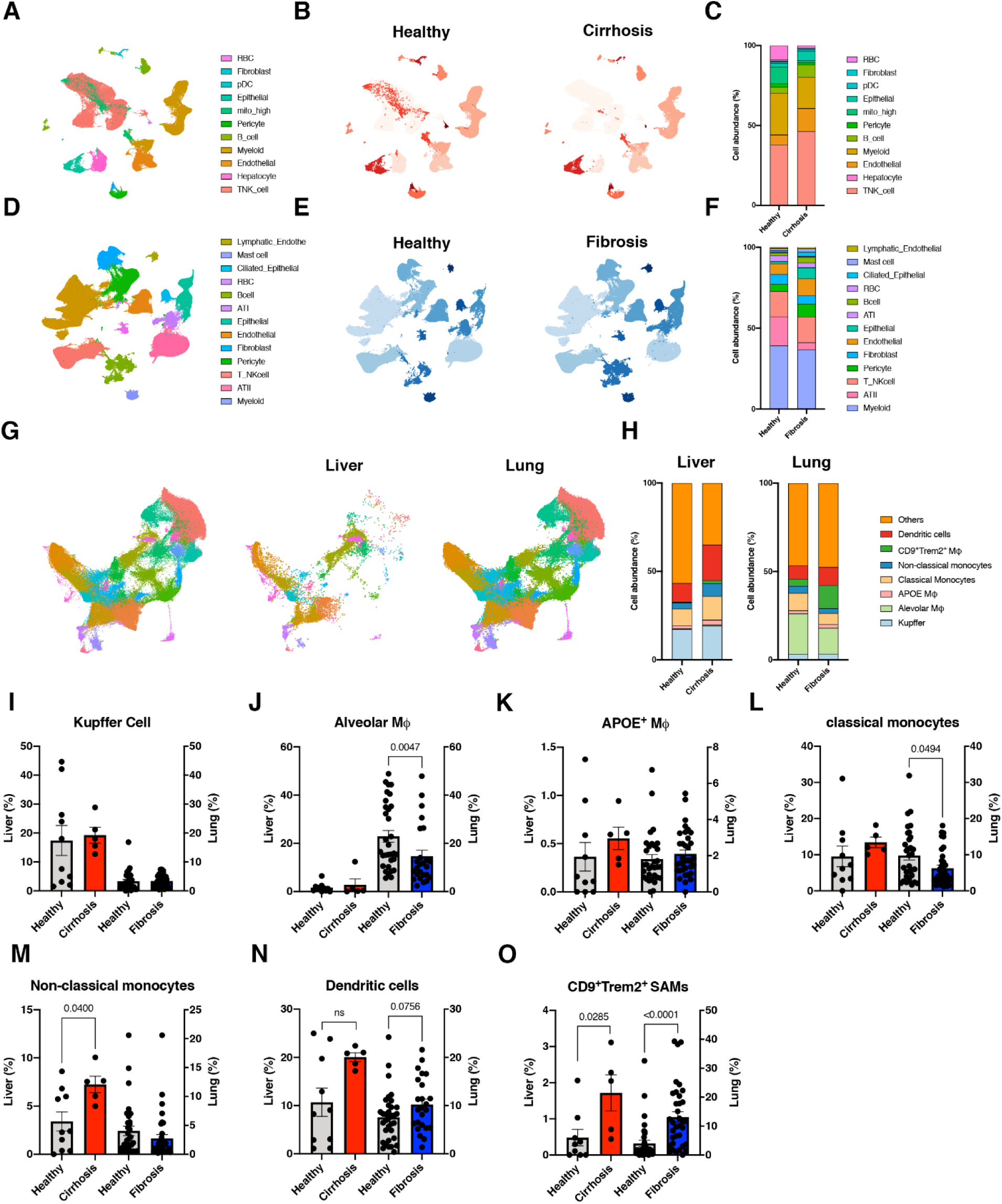
Integration of human single cell RNA-seq lung and liver fibrosis datasets identified *CD9*^+^*TREM2*^+^ scar-associated macrophages as core fibrosis-associated subsets. (A) Uniform manifold approximation and projection (UMAP) plot of all cells from integrated liver datasets colored by cell type. (B) UMAP plots of cells from healthy and cirrhotic livers shaded by cell type. (C) Stacked bar chart of cell subset frequencies from liver. (D) UMAP plot of all cells from integrated lung datasets colored by cell type. (E) UMAP plots of cells from healthy and fibrotic lungs shaded by cell type. (F) Stacked bar chart of cell subset frequencies from lung. (G) UMAP plot of myeloid atlas from integrated lung and liver datasets colored by cell type. (H) Stacked bar chart of simplified myeloid cell subset frequencies from liver and lungs. Frequency per sample of Kupffer cells (I), alveolar macrophages (J), *APOE*^+^ macrophages (K), classical monocytes (L), non-classical monocytes (M), and dendritic cells (N). (O) Increased frequency of *CD9*^+^*TREM2*^+^ scar-associated macrophages in cirrhotic livers (red) and fibrotic lungs (blue). Statistics Mann-Whitney u test, p-values displayed on plots.

**Figure 2.**
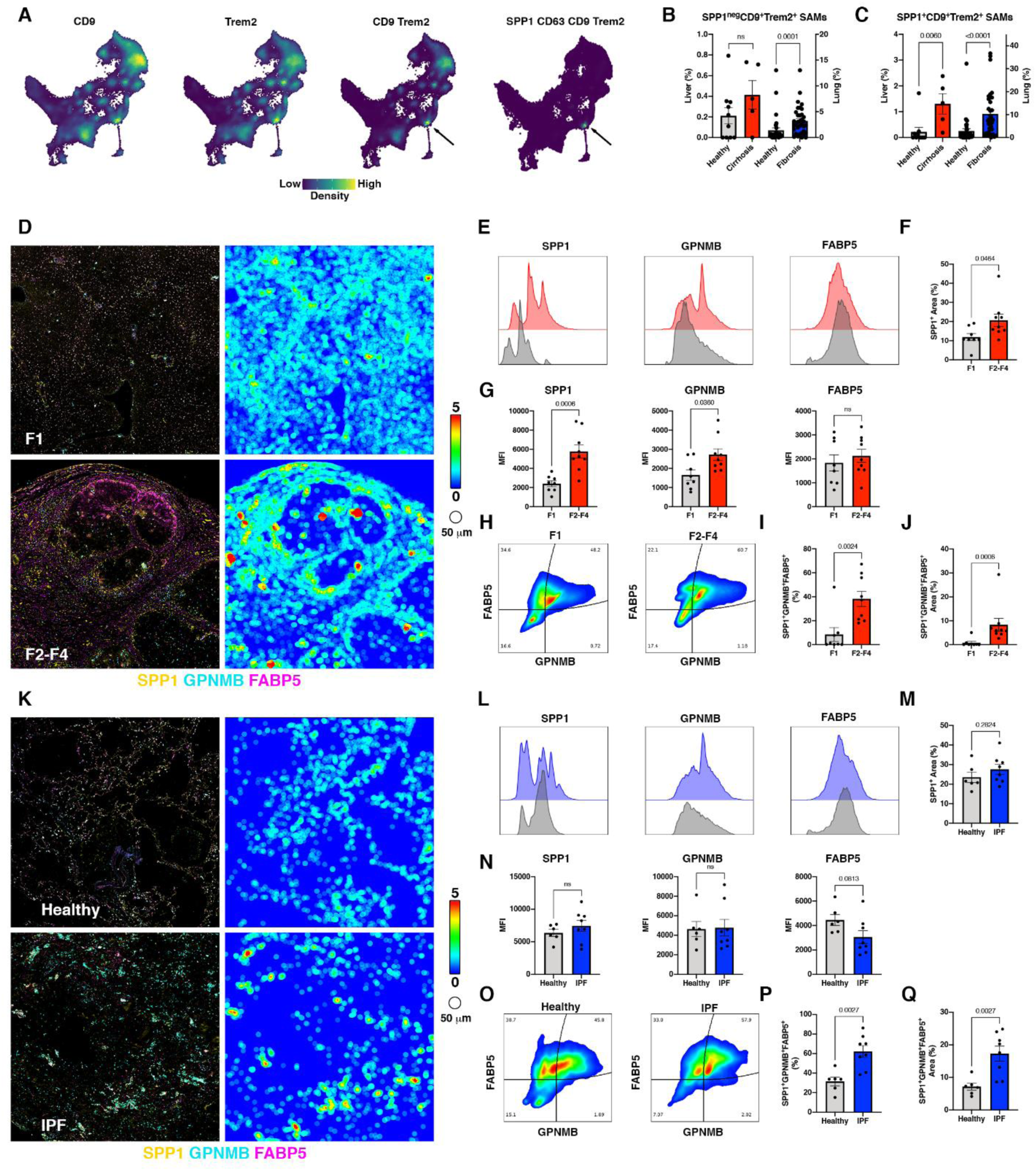
CD9^+^TREM2^+^ macrophages in human liver and lung fibrotic disease are a heterogenous population, from which accumulating scar associated macrophages can be further resolved via co-expression of SPP1, GPNMB and FABP5. (A) Nebulosa plot showing density of CD9, TREM2, joint density of CD9 and TREM2, or joint density of SPP1, CD63, CD9 and TREM2 allowing precise identification of scar-associated macrophages. (B-C) Frequency of SPP1^neg^ (B) and SPP1^+^ (C) scar-associated macrophages in liver and lung. (D and K) CycIF of SPP1, GPNMB and FABP5 in human F1 and F2-F4 non-alcoholic steatohepatitis (NASH) (D) and healthy and idiopathic pulmonary fibrosis (IPF) (K). Representative tissue heatmaps (scale, blue = 0 to red = 5 cells/50 μm) of SPP1^+^ cells. (E and L) Histogram representation of SPP1, GPNMB and FAPB5 median fluorescence intensity (MFI)/cell in liver (E) and lung (L). (F and M) SPP1^+^ area in liver (F) and lung (M). (G and N) Average expression per cell of SPP1, GPNMB and FAPB5 in liver (G) and lung (N). (H and O) Representative plot of SPP1^+^ cells co-expressing GPNMB and FABP5 in liver (H) and lung (O). (I and P) Increased frequency of SPP1^+^GPNMB^+^FABP5^+^ cells in fibrotic livers (I) and lungs (P). (J and Q) Increased area of SPP1^+^ scar-associated macrophages in fibrotic livers (J) and lungs (Q). Statistics Mann-Whitney u test, p-values displayed on plots.

**Figure 3.**
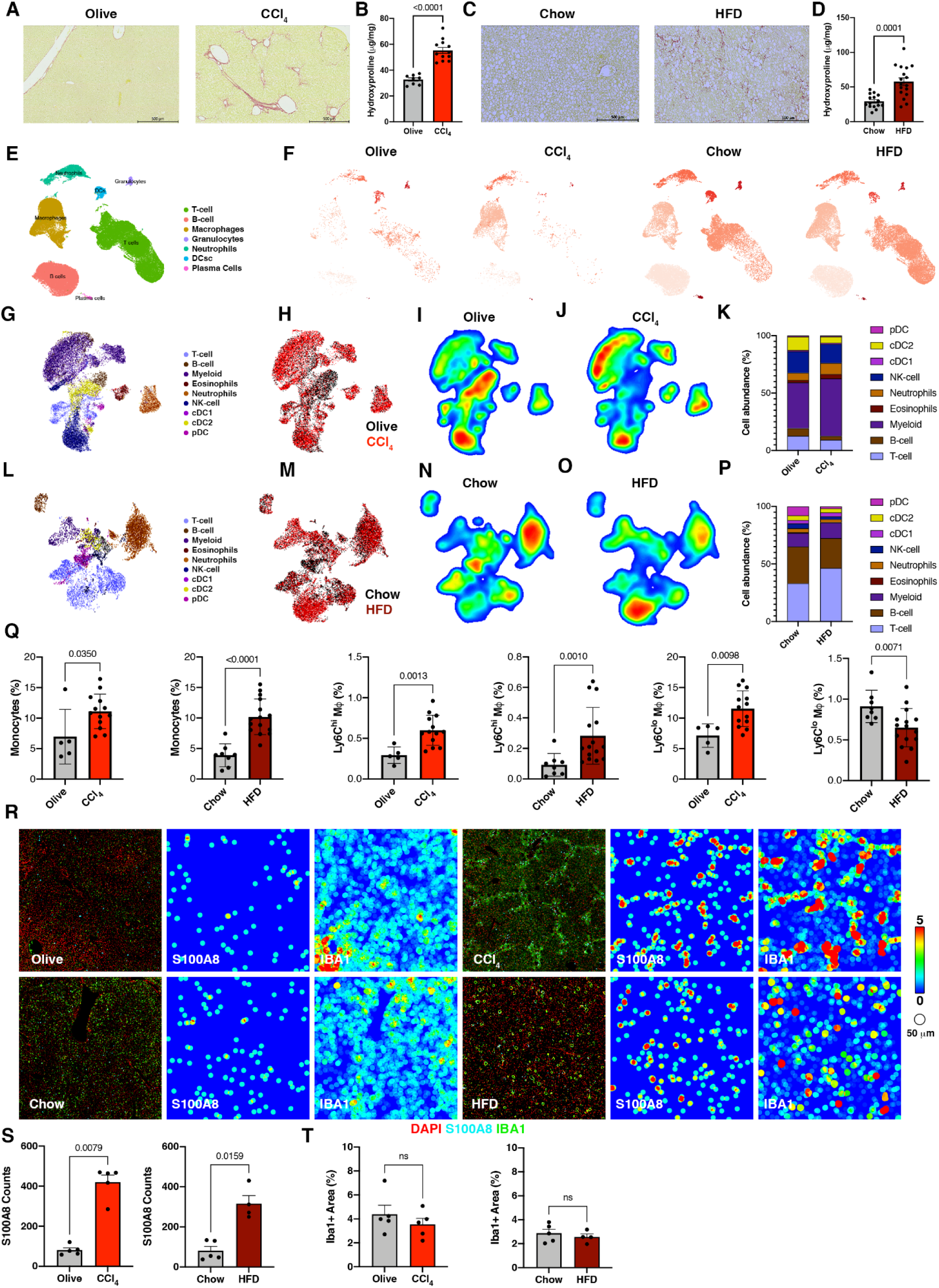
Myeloid cells and neutrophils accumulate within and at the edge of the scar in murine liver fibrosis models. (A) Representative picro Sirius red staining of olive oil and CCl_4_ livers. (B) Quantification of hydroxyproline content in CCl_4_ injury. (C) Representative images of picro Sirius Red staining of chow and high fat diet (HFD) livers. (D) Quantification of hydroxyproline content in HFD. (E) UMAP single-cell RNA-seq plot of all cells from olive-oil/CCl_4_- and chow/HFD-treated livers. (F) UMAP single-cell RNA-seq plots of liver cells split by condition. (G and H) UMAP FACS plot of CD45^+^ cells from CCl_4_ model colored by cell type (G) or by condition (olive oil (black) and CCl_4_ (red)). (I and J) UMAP FACS density plots of CD45^+^ cells in olive oil (I) and CCl_4_ (J) showing increased frequency of myeloid cells. (K) Stacked bar chart of CD45^+^ cell populations in CCl_4_ injury. (L and M) UMAP FACS plots of CD45^+^ cells from HFD model colored by cell type (L) or condition (chow (black) and HFD (dark red)) (M). (N and O) UMAP FACS density plots of CD45^+^ cells in chow (N) and HFD (O) showing increased frequency of myeloid cells. (P) Stacked bar chart of CD45^+^ cell populations in HFD. (Q) Frequency of Ly6C^hi^ monocytes, Ly6C^hi^CD68^+^F4/80^+^ macrophages and Ly6C^lo^CD68^+^F4/80^+^ macrophages in CCl_4_ (red) and HFD (dark red). (R) CycIF of S100A8 (neutrophils) and Iba1 (macrophages) in CCl_4_ and HFD. Representative tissue heatmaps (scale, blue = 0 to red = 5 cells/50 μm) of S100A8 and Iba1 showing accumulation in the scar. (S) Increased numbers of neutrophils in CCl_4_ and HFD models. (T) Iba1^+^ positive area. Statistics Mann-Whitney u test, p-values displayed on plots.

For murine samples, expression FASTQs were pre-processed with Cell Ranger (10x Genomics) using the default settings and aligned to the provided mm10 reference genome (Ensembl 93, 10x Genomics, (*33*)). These were further pre-processed through the velocyto package using the run10x wrapper, mm10 genome above and mm10 repeat mask (UCSC Genome Browser, https://genome.ucsc.edu/index.html, (*34, 35*)). Count matrices for gene expression, hashing barcodes, spliced, unspliced and ambiguous reads were combined into a single Seurat Object for each sample or sequencing lane (Seurat version 4.0.3 (*36*), R version 4.0.0, RStudio version 1.2.5042). Lipid hashed samples were demultiplexed as described by McGinnis *et al.* (*28*) or using HTODemux in Seurat (*36, 37*). Aligned, demultiplexed matrices from all experiments were combined and barcodes with greater than 25% mitochondrial reads and obvious multiplets (clusters identified by known marker genes of at least two cell types) were removed. Genes per cell, transcript counts per cell and mitochondrial read percentage per cell are presented in Supplemental Figure 6.

#### Cell Clustering & Annotation

Seurat was used for dimensionality reduction, cell clustering and differential gene analysis for both human and murine samples. Human single cell objects were split by dataset and chemistry and normalized using SCTransform with percent mitochondrial reads regressed out (*38*). The top 1000 variable features from each dataset were combined and used in downstream steps. Harmony (*39*) was used to correct for patient, dataset, and chemistry variability and 30 PCs were used for RunUMAP and FindNeighbors. A resolution of 0.7 was used for cluster selection. FindAllMarkers was used to identify markers for clusters and annotation was performed manually using the markers shown in Supplemental Figures 1, 2 and 3 (for the human liver, lung and myeloid atlases, respectively).

Murine single cell objects were split by experiment and normalized as above. The top 3000 variable features were combined and used in downstream steps. Integration was performed across experiments using Harmony (*39*). 16 PCs were used for RunUMAP and FindNeighbors, and cluster selection was performed using a resolution of 0.1 based on ClusTree stability diagrams (*40*). Clusters were manually annotated using the markers shown in Supplemental Figure 4. For high-resolution analysis of monocyte and macrophage populations, this subset was extracted, re-normalized and re-integrated as above using 3000 variable features. 20 PCs were used for RunUMAP and FindNeighbors. Cluster markers were identified with FindAllMarkers, and clusters were manually annotated by comparing cluster marker lists to prior publications and the human dataset presented in this paper ((*5, 12, 14, 17-19, 21, 26*) and Supplemental Figure 6.

#### RNA Velocity & Pseudotime Ordering of Monocytes & Macrophages

Spliced and unspliced read counts were estimated as described in Data Preprocessing & Quality Control. RNA velocity was modeled using the scVelo package (*35, 41*). Briefly, the myeloid compartment counts were normalized, and cells were clustered and annotated as previously described using Seurat. The cells were separated into their respective treatment groups and velocity was calculated using the functions tl.recover_dynamics and tl.velocity from the scVelo package (default settings and mode = ‘dynamical’).

### Tissue processing

Murine liver and lung were collected in RPMI1640 media on ice, minced and digested in 100 U/mL collagenase (Sigma-Aldrich) at 37°C for 1 hour with shaking. Digested tissue was filtered through 70 μm nylon mesh and hepatocytes removed by spinning at 50x*g* for 3 minutes. The remaining cells were passed through a 25 % Percoll (GE Life Sciences) gradient and erythrocytes were lysed with ammonium-chloride-potassium buffer. Single cells were aliquoted for flow cytometry or pooled and distributed for restimulation and scRNAseq.

### Multiplex cytokine analysis

Intra-hepatic leukocytes (IHLs) or intra-pulmonary leukocytes (IPLs) were resuspended at a concentration of 5 million cells per mL in RPMI with 10% FBS and 1X stimulation cocktail (ebioscience) for 18 hours. Supernatant were collected and frozen prior to cytokine quantification. Multiplex cytokine or chemokine quantification was performed on 25 μl of supertanant using Legendplex (Biolegend). Analysis was performed using Biolegend analysis software. Data were normalized to naïve animals and expressed as fold-change.

### Flow Cytometry

Murine and human cells were washed with PBS and stained with LIVE/DEAD fixable aqua viability dye (diluted 1:250 in PBS; Invitrogen) for 20 minutes at room temperature. Following washing with FACS buffer (PBS supplemented with 2% fetal bovine serum (FBS), 10 mM HEPES and 2 mM EDTA) cells were fixed with Foxp3/Transcription Factor Fixation/Permeabilization buffer (Invitrogen) for 30 minutes at 4°C. After washing cells were resuspended in 90% FACS buffer/10% DMSO and stored at -80°C until antibody staining, and flow cytometry were performed.

After thawing cells were washed and Fc receptors were blocked with TruStain FcX (mouse or human as appropriate; Biolegend) for 5 minutes at room temperature before staining with diluted antibodies (Supplemental Tables 2, 3 and 4). Cells were then washed, filtered through 60 μm mesh and run on an LSRFortessa (BD). Data were analyzed in FlowJo version 10.5.+ (BD) using the DownSample and UMAP plugins.

### Cyclic Immunofluorescence staining

Cyclic multiplex immunofluorescence (cycIF) was performed as described in Molina *et al.* using the antibodies described in Supplemental Tables 5 (for human sections) and 6 (for murine sections) (*42*). Briefly, formalin-fixed, paraffin-embedded (FFPE) sections of human or murine liver or lung specimens were deparaffinized, rehydrated and subjected to heat-induced epitope retrieval. After blocking with normal donkey serum slides were stained with primary antibodies overnight. Unbound antibodies were removed by washing and slides were stained with fluorescently-conjugated secondary antibodies made in donkeys (Supplemental Table 7). Slides were again washed, DNA stained with DAPI (Invitrogen) and mounted in ProLong Diamond Antifade Mountant with DAPI (Invitrogen). Stained slides were imaged using an Axio Scan (Zeiss). Following each round of imaging, coverslips were removed and antibodies stripped.

### Histology

Deparaffinization of FFPE sections was performed with two 5 minutes Xylene and 100% Ethanol bath. Sections were then incubated in Picro Sirius Red solution (Direct Red 81 0.2% in 1.3% Picric Acid Solution, Sigma) for 1 hour. Sections were rinsed in dH2O, then Xylene:Ethanol and dried in Xylene for 5 minutes. Sections were mounted with Permount (Fisher) and acquired on a Leica system. Quantification of red pixels was then performed to quantify percent area of fibrosis.

### Image Analysis

Acquired images were first subjected to shading correction, as well as stitching using the Zeiss ZEN blue DESK (version 3.0) image processing software. The alignment across subsequent rounds was performed on their respective nuclear (DAPI) channel *via* FIJI and applying the transformation to the remaining channels (*43, 44*). The multi-channel image pyramid was generated using Bioformats2Raw and Raw2OMETiff programs (both Glencoe Software, Seattle, WA) and finally loaded into Visiopharm (Visiopharm A/S, Denmark) for analyses. A representative 2 mm x 2 mm ROI was centered on the tissue, where segmentation and heatmap APPs were applied for individual markers. All APPs were run with the same parameters across the experimental groups within the same tissue type to ensure data integrity. Numerical values were extracted and processed using Microsoft Excel and R and converted to a FlowJo-compatible format (FCS) for further analyses.

### Cell Culture (Macs in vitro/co-culture)

Cryopreserved primary human CD14+ monocytes (Stemcell Technologies) isolated from peripheral blood were plated at 300,000 cells/well in flat bottom, tissue culture-treated 96-well plates (BD Falcon) in RPMI 1640 (Gibco) supplemented with 10% fetal bovine serum (Gibco), 10 mM HEPES (Gibco) and 2 mM GlutaMax (Gibco). Monocytes were cultured in 10 ng/mL recombinant human GM-CSF or M-CSF (both Biolegend) for 7 days at 37°C, 5% CO_2_ in normoxic conditions. Media and cytokines were refreshed on day 3 of culture. Conditioned media was frozen at -80°C. Macrophages were washed with PBS and detached by incubating in cold PBS at 4°C for at least 30 minutes followed by vigorous pipetting and scraping.

### High content imaging

For monocyte/macrophage monocultures cryopreserved primary human CD14^+^ monocytes (Stemcell Technologies) isolated from peripheral blood were plated at 300,000 cells/well in flat bottom, tissue culture-treated 96-well plates (BD Falcon) in complete RPMI (RPMI 1640 (Gibco) supplemented with 10% fetal bovine serum (Gibco), 10 mM HEPES (Gibco), 100 U/mL each of penicillin and streptomycin (Gibco) and 2mM GlutaMax (Gibco)). Monocytes were cultured in 10 ng/mL recombinant human GM-CSF or M-CSF (both Biolegend) for 7 days at 37°C, 5% CO_2_ in normoxic conditions. Media and cytokines were refreshed on day 3 of culture. Conditioned media was frozen at -80°C. Macrophages were washed with PBS and detached by incubating in cold PBS at 4°C for at least 30 minutes followed by vigorous pipetting and scraping.

Primary human hepatic stellate cells (HSC; used between passages 4 and 8) and primary human GM-CSF monocyte derived macrophages (MDM) were cultured in collagen-1-coated CellCarrier Ultra 96-well black walled plates (PerkinElmer) at a ratio of 1425 HSC:28500 MDM in FBM (Lonza) supplemented with 0.2% fetal bovine serum (FBS; Gibco) and 300 μM L-ascorbic acid 2-phosphate (Sigma). Recombinant human interferon gamma (IFNγ; Biolegend) and recombinant human transforming growth factor beta-1 (TGF-β1; Biolegend) were used at final concentrations of 10 ng/mL. Co-cultures were incubated for 7 days at 37°C, 5% CO_2_ in normoxic conditions without changing the media or refreshing cytokines. Cells were fixed with cold 95% methanol/5% glacial acetic acid (Sigma) and washed extensively with PBS. Blocking was performed with 5% normal goat serum (Gibco) followed by staining with mouse IgG2a anti-human α- Smooth muscle actin (SMA; clone 1A4; Sigma) and mouse IgG1 anti-Collagen 1 (clone Col1; Sigma). Primary antibodies were detected with goat anti-mouse IgG2a AlexaFluor 647 and goat anti-mouse IgG1 AlexaFluor 568 (both Invitrogen). Cytoplasm was detected with HCS CellMask Green (Invitrogen) and nuclei with Hoechst 33342 (BD Biosciences). Imaging was performed on an Opera Phenix (PerkinElmer) with a 10x objective. Total Collagen-1 intensity is presented without normalizing to cell number because macrophages plated alone did not produce detectable Collagen-1.

### Statistical Analysis

All statistical tests were conducted using R version 4.0.0 (https://www.r-project.org/) or Prism version 9. Differential expression testing was performed in R using Seurat version 4.0.0. Graphs were generated using GraphPad Prism 9 (GraphPad Software, La Jolla, CA), the R (version 4.0.0) packages Seurat version 4.0.0 (*36*), ggplot2 version 3.3.5 and Nebulosa version 0.99.92 (*45*) in RStudio (version 1.2.5042), Visiopharm or FlowJo. Differences between two groups were determined by Mann-Whitney, whereas differences between groups were determined by ANOVA followed by Tukey post-hoc test. Kruskal–Wallis tests was used when group sizes were too different and if the data did not data meet the assumption of homogeneity of variance.

### Results

#### Integration of human single cell (sc) RNAseq datasets identifies a core scar-associated macrophage subset

To better characterize macrophage populations in fibrosis, we integrated publicly available single cell RNA sequencing data from liver and lung fibrosis patients. This demonstrated dysregulation in multiple leukocyte populations including myeloid cells (Fig. 1A-F). To better characterize macrophage subsets associated with fibrosis, we created a myeloid cell atlas from these combined data (Fig. 1G-H). We confirmed the biological validity of our integration by demonstrating that Kupffer cells are restricted to the liver data, alveolar macrophages are restricted to the lung data (Fig. 1I-J). Several myeloid populations demonstrated organ-specific changes in fibrosis (Fig. 1K-N). Our myeloid atlas approach similarly recapitulated the previous finding that *TREM2*^+^*CD9*^+^ scar-associated macrophages expand in human liver fibrosis, and we now reveal that a similar population is also expanded in human lung fibrosis (Fig. 1O).

#### SPP1, GPNMB and FABP5 resolve the scar-associated macrophages from the heterogeneous CD9^+^TREM2^+^ population in human fibrosis

Examining the expression of *CD9* and *TREM2* within our myeloid atlas demonstrated that this combination of markers identifies multiple populations in human liver and lung fibrosis (Fig. 2A and Sup. Fig. 4A). This indicated granularity within the *CD9*^+^*TREM2*^+^ population that was better resolved upon the addition of *SPP1* and *CD63*. For example, there was no increase in *CD9*^+^*TREM2*^+^*SPP1*^-^ macrophages in liver fibrosis, and a slight increase in lung fibrosis (from 2% to 4%; 2-fold increase, p=0.001); however, *CD9*^+^*TREM2*^+^*SPP1*^+^ macrophages had a more dramatic increase in both liver (0.2% to1.3 %; 6.5-fold increase, p=0.006) and lung fibrosis (2% to 10%; 5-fold increase, p<0.0001) (Fig. 2B-C). We confirmed that these CD9^+^TREM2^+^SPP1^+^ macrophages were indeed scar-associated by performing immunofluorescent staining in fibrotic human NASH liver and IPF lung sections, which also allowed us to validate the increases in these markers at the protein level (Fig. 2D-Q). Curating the data, we identified that the combination of SPP1, GPNMB and FABP5 had the best specificity for identifying these scar-associated macrophages, which was independently confirmed by the unbiased tool COMET (Fig. 2A and Sup. Fig. 4B). Herein we refer to this as the SPP1 scar-associated signature. We then demonstrated increased density of SPP1 in cirrhotic F4 vs F1 NASH livers (Fig. 2D, F). Interestingly, there was a significant increase in the expression of SPP1 and GPNMB, but not in FABP5, per cell in F4 patients (Fig. 2G). Furthermore, there was an increase in frequency of triple SPP1, GPNMB and FABP5 positive scar-associated macrophages in both cirrhotic NASH livers and IPF lungs (Fig. 2H-I) with a concomitant increase in the area covered by these triple-positive cells (Fig. 2J and Q). Despite increased SPP1 density in IPF lung (Fig. 2K), there was no increase in the frequency or area of single-positive cells (Fig. 2L-N). Combining these three markers was strictly required to identify the expansion in frequency and area covered by scar-associated macrophages in IPF lungs (Fig. 2O-Q). Interestingly, scar-associated macrophages were found at the edge of the scar at the interface between pathogenic fibrillar- and normal extracellular matrix (ECM) (Sup. Fig. 5). Thus, this analysis uncovered that a subpopulation of CD9^+^TREM2^+^SPP1^+^, GPNMB^+^ and FABP5^+^ macrophages (designated “Fab5” or five marker positive SAMs) are the population that is most increased and scar-associated in human liver and lung fibrosis.

#### Myeloid cells and neutrophils accumulate within and at the edge of the scar in pre-clinical liver fibrosis models

To extend these observations to animal models of fibrosis, we performed scRNAseq on livers from mice treated with carbon tetrachloride (CCl_4_) or fed high-fat diet (HFD). Prior to investigating whether there was a similar scar-associated macrophage population in these models, we first confirmed that these mice developed robust fibrosis (Fig. 3A-D). Our scRNAseq demonstrated dysregulation of multiple leukocyte populations, including a significant alteration in myeloid populations (Fig. 3E-F). Through flow cytometry analysis we confirmed the dysregulation seen by scRNAseq (Fig. 3G-P). By flow, we observed an increase in Ly6C^hi^ monocytes and macrophages in both CCl_4_ and HFD, and an increase in Ly6C^lo^ macrophages in CCl_4_ but decrease in HFD (Fig. 3Q). Importantly, we confirmed a reorganization of Iba1^+^ macrophages aligning with scarred areas and increased S100A8/A9^+^ neutrophils that co-localized with these macrophages in the scar (Fig. 3R-T).

#### Fab5 macrophages originate from monocytes based on their transcriptional trajectories and specifically accumulate in fibrotic lesions

Having recapitulated the fibrosis-associated macrophage dysregulation seen in patients in murine models of liver fibrosis, we subclustered the macrophages and identified the same scRNAseq Spp1^+^ signature (Fig. 4A and Sup. Fig. 7). Applying RNA Velocity suggested that monocytes pass through an inflammatory macrophage state before acquiring the Fab5 signature as seen previously (Fig. 4B-D) (*11, 12, 14*). To confirm that this population was associated with fibrosis progression, we performed flow cytometry using a surrogate gating strategy due to lack of flow reagents for murine SPP1 and FABP5. Manual curation and COMET demonstrated that the combination of *Trem2*, *Cd9*, *Cd63* and *Gpnmb* was equally selective for this population (Sup. Fig. 8). This combination of markers validated the fibrosis-associated increase in CD9^+^TREM2^+^CD63^hi^ macrophages (Fig. 4E-G). Critically, cycIF using the same gating strategy (SPP1^+^GPNMB^+^FABP5^+^) employed in patient samples validated the increase and physical proximity of these macrophages to the scar (Fig. 4H). This analysis also recapitulated the increased density of SPP1 and triple-positive macrophages seen in human cirrhotic livers (Fig. 4I-N). Similar analyses performed in the bleomycin model of lung fibrosis supported this being a core fibrogenic signature as in the human data (Sup. Fig. 9).

**Figure 4.**
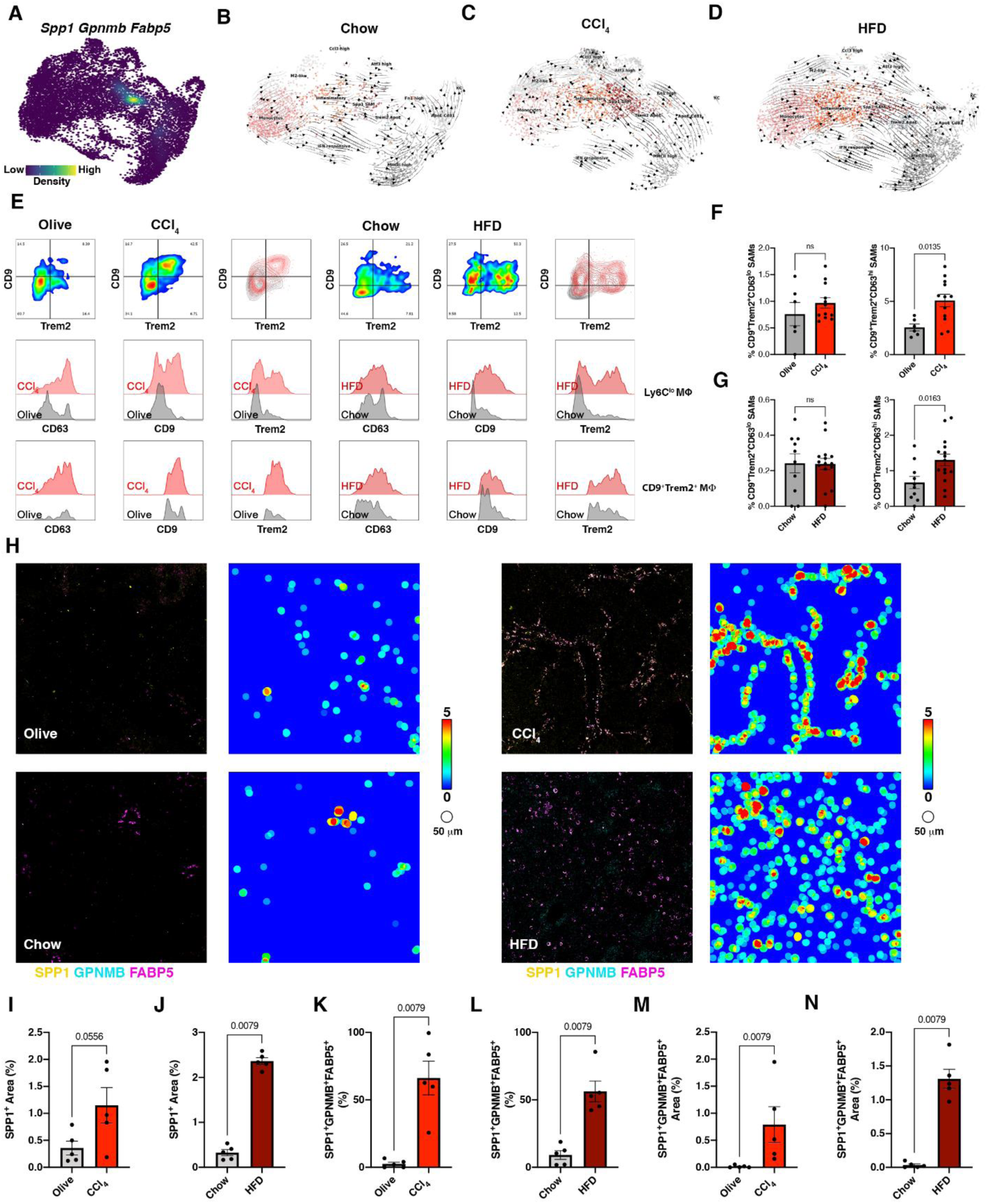
SPP1^+^ scar-associated macrophages originate from monocytes based on their transcriptional trajectories, and specifically accumulate in fibrotic lesions in two murine liver fibrosis models. (A) Nebulosa plot showing joint density of scar-associated macrophages genes *Spp1*, *Gpnmb* and *Fabp5*. (B-D) RNA velocity field projected onto UMAP plots of myeloid cells from chow (B), CCl_4_ (C) and HFD (D) livers. (E) Representative FACS plots of CD9 and Trem2 on Ly6C^lo^F4/80^+^ hepatic macrophages from olive (grey), CCl_4_ (red), chow (grey) and HFD (dark red). Histograms of CD63, CD9 and Trem2 on Ly6C^lo^F4/80^+^ hepatic macrophages and CD9^+^Trem2^+^ scar-associated macrophages. (F and G) Frequency of CD9^+^Trem2^+^CD63^lo^ (right) and CD9^+^Trem2^+^CD63^hi^ (left) scar-associated macrophages in CCl_4_ (F) and HFD (G). (H) CycIF staining of SPP1, GPNMB and FAPB5 from Olive, CCl_4_, chow and HFD livers. Representative tissue heatmaps (scale, blue = 0 to red = 5 cells/50 μm) of SPP1^+^ cells. (I and J) Total SPP1-positive area in CCl_4_ (I) and HFD (J). (K and L) Frequency of SPP1^+^GPNMB^+^FABP5^+^ scar-associated macrophages in CCl_4_ (K) and HFD (L). (M and N) Area of SPP1^+^GPNMB^+^FABP5^+^ scar-associated macrophages in CCl_4_ (M) and HFD (N). Statistics Mann-Whitney u test, p-values displayed on plots.

#### In vitro drivers of Fab5 macrophage differentiation and functions

We next investigated cytokines that could drive the Fab5 signature in human monocytes. First, we confirmed mixed type 1, 2 and 3 cytokine production by murine intrahepatic lymphocytes from CCl_4_ and HFD animals stimulated *ex vivo* (Fig. 5A-B). Using cycIF we confirmed that neutrophils producing type 3 cytokines IL-17A and GM-CSF physically co-localize with macrophages within the hepatic scar (Fig. 5C-I). Furthermore, we found that TGF-β-activating molecules like CD29 and MMP9 were similarly enriched and co-located in the scar (Fig. 5C-I). Based on the physical proximity between GM-CSF-producing neutrophils and macrophages in the scar, we hypothesized that GM-CSF would drive monocytes towards this signature *in vitro*. Comparison of M-CSF and GM-CSF differentiation confirmed this hypothesis (Fig. 5J-K). We then investigated the roles of the main type 1, 2 and 3 cytokines. Type 1 (IFNγ) and 2 (IL-4/IL-13) cytokines inhibited this phenotype, as seen by a decreased frequency of CD9, TREM2 and CD63 (Fig. 5L-N). Conversely, the type 3 cytokines IL-17A and TGF-β1 increased the frequency of macrophages expressing this Fab5 signature (Fig. 5L-N).

**Figure 5.**
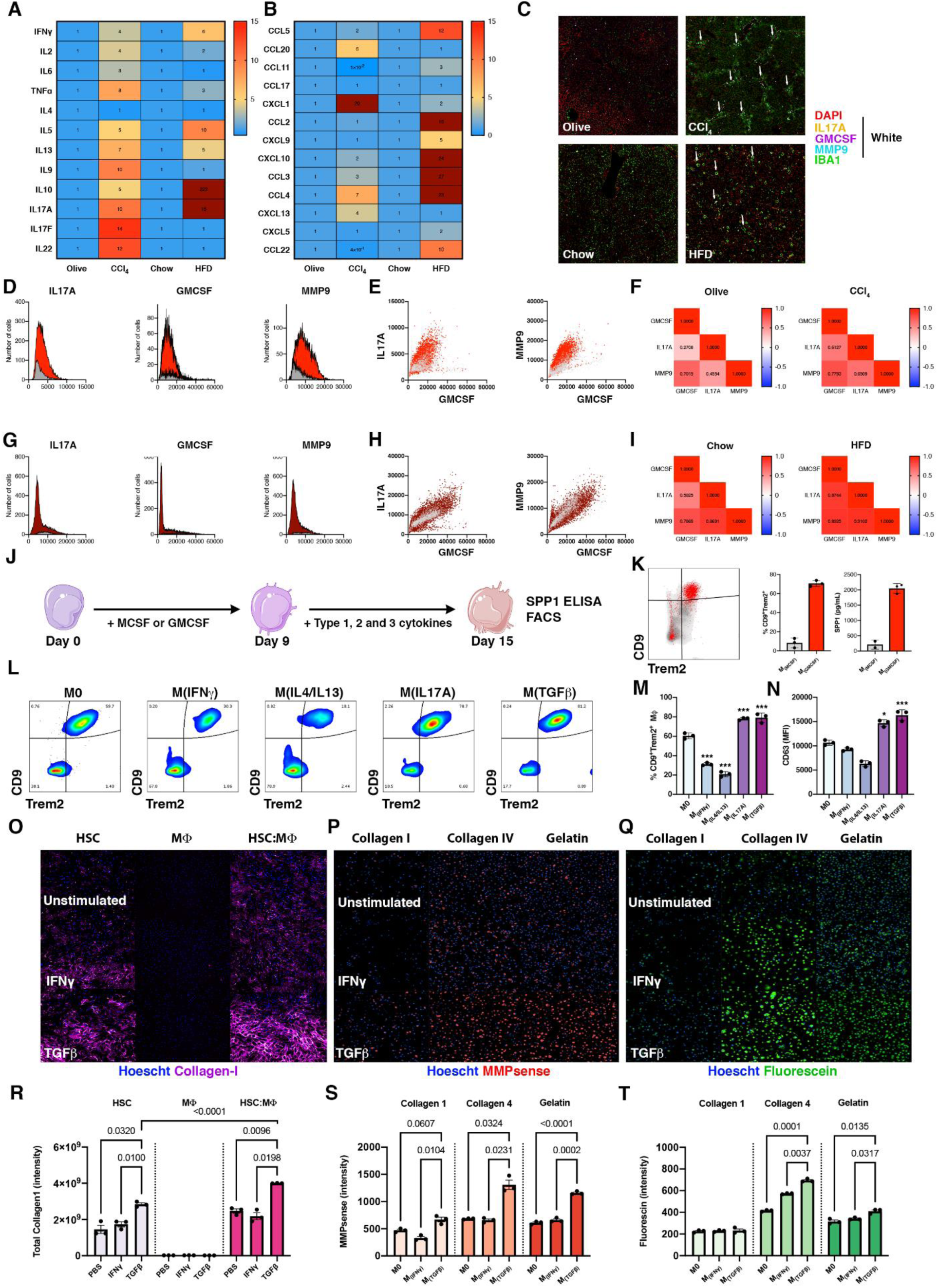
TGF-β, IL-17A and GM-CSF are sufficient to induce phenotypic differentiation of SPP1^+^ scar-associated macrophages from human monocytes. (A and B) Heatmap comparing fold change in type 1, 2 and 3 cytokines (A) and chemokines (B) from stimulated intra-hepatic lymphocytes from CCl_4_ and HFD. (C) CycIF staining of Iba1, IL-17A, GM-CSF and MMP9 showing localization of type 3 cytokines. (D) Histogram of IL-17A, GM-CSF and MMP9 expression by S100A8^+^ neutrophils in olive oil (grey) and CCl_4_ (red) livers. (E) Correlation plots between IL-17A, GM-CSF and MMP9 in olive oil (grey) and CCl_4_ (red). (F) Correlation heatmap of IL-17A, GM-CSF and MMP9 in olive oil and CCl_4_ livers. (G) Histogram of IL-17A, GM-CSF and MMP9 expression by S100A8^+^ neutrophils in chow (grey) and HFD (dark red) livers. (I) Correlation plots between IL-17A, GM-CSF and MMP9 in chow (grey) and HFD (dark red) livers. (J) Schematic of in vitro macrophage differentiation assay. (K) Representative CD9 vs. Trem2 FACS plot of M-CSF-derived (grey) or GM-CSF-derived (red) macrophages at day 9. Frequency of CD9^+^Trem2^+^ macrophages (left) and SPP1 concentration in supernatant (right) at day 15. (L) Representative CD9 vs. Trem2 FACS plot of GM-CSF-derived M0, M(IFNγ), M(IL4/IL13), M(IL-17A) and M(TGF-β). (M) Frequency of CD9^+^Trem2^+^ macrophages following cytokine stimulations. (N) CD63 MFI of GMCSF-derived M0, M(IFNγ), M(IL4/IL13), M(IL-17A) and M(TGF-β). (O) Collagen 1 staining (purple) in co-cultures of macrophages with primary human hepatic stellate cells stimulated with either IFNγ or TGF-β1. (R) Quantification of collagen 1 average intensity per cell in HSC mono-culture (light purple), macrophage mono-culture (pink) and HSC:macrophage co-culture (Purple). (P) MMPsense (red) or DQ-fluorecein (Q) (Green) staining of macrophages cultured on either DQ-Collagen I, DQ-Collagen IV or DQ-Gelatin substrates stimulated with either IFNγ or TGF-β1. (S) Quantification of MMPsense per cell and Fluorescein per well (T) from macrophages cultured on either DQ-Collagen I (light color), DQ-Collagen IV (color) or DQ-Gelatin (dark color) substrates stimulated with either IFNγ or TGF-β1. Statistics ANOVA or 2-way ANOVA followed by a post hoc Tukey, p-values displayed on plots.

To confirm the pro-fibrotic role of these macrophages we co-cultured primary human macrophages and hepatic stellate cells (HSC) with pro-inflammatory IFNγ or pro-fibrotic TGF-β1. As expected, TGF-β1 enhanced type 1 collagen production by co-cultured HSC correlating with induction of the Fab5 signature (Fig. 5O and R). GOrilla pathway analysis of the *SPP1* SAM marker genes demonstrated an enrichment for extracellular matrix production-associated transcripts (data not shown). We next investigated if *in vitro* differentiated scar-associated macrophages (M_TGFβ_) showed differential ECM-remodeling activity. GM-CSF derived macrophages (M0) stimulated with either IFNγ (M_IFNγ_) or TGF-β (M_TGFβ_) were cultured on DQ-COL I (scar), DQ-COL IV or DQ-Gelatin (both normal ECM) substrates. Interestingly, M_TGFβ_ demonstrated higher MMP activity when compared to M0 and M_IFNγ_ which was significantly upregulated on normal ECM (Fig. 5P and S). This phenotype correlated with increased degradation of normal ECM (Fig. 5Q and T). Together these data indicate that GM-CSF, IL-17A and TGF-β collaborate to induce monocyte-to-SAM differentiation, and suggest that SAM both degrade normal ECM and promote pathogenic collagen production by mesenchymal cells.

#### Therapeutic TGF-β, IL-17A or GM-CSF blockade reduced Fab5 macrophages and fibrosis in chronic CCl_4_ injury

To validate our *in vitro* findings suggesting that TGF-β, IL-17A and GM-CSF are important drivers of both scar-associated macrophages and fibrosis, we therapeutically inhibited each of these three cytokines in chronic CCl_4_ liver injury. We observed a significant reduction of fibrosis measured by histology and hydroxyproline liver content in all treatment groups (Fig. 6A-B). As shown previously, fibrosis progression was associated with an increased frequency of myeloid cells by flow cytometry (Fig. 6C-D). Interestingly, neutralization of IL-17A or GM-CSF led to a reduced frequency of CD11b^+^ cells out of CD45^+^ cells infiltrating the liver in contrast to TGF-β blockade (Fig. 6D-E). TGF-β inhibition significantly decreased the frequency of Ly6C^lo^ macrophages, CD63^lo^ scar-associated macrophages and CD63^hi^ scar-associated macrophages (Fig. 6F, L-O). In contrast, IL-17A and GM-CSF blockades only impacted the frequency of CD63^hi^ scar-associated macrophages (Fig. 6L-O). All treatments also reduced the levels of CD63 and GPNMB (Fig. 6J-K) expressed by these cells reinforcing our *in vitro* observation (Fig. 5L-N). These data validate the importance of TGF-β, IL-17A and GM-CSF in driving scar-associated macrophage differentiation *in vivo* and further support the connection between Fab5 macrophages and fibrosis severity.

**Figure 6.**
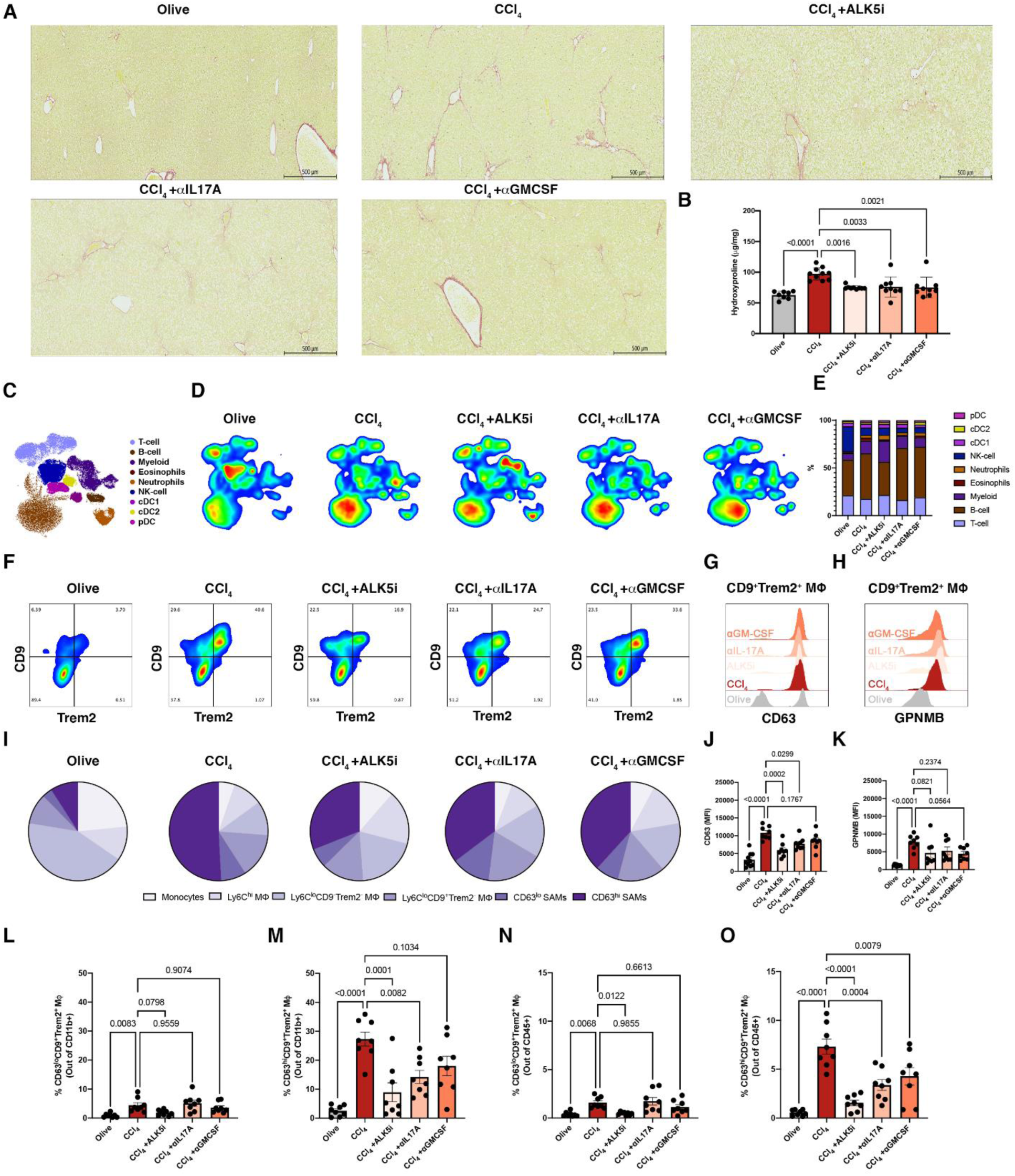
Therapeutic TGF-β, IL-17A or GM-CSF blockade inhibits the differentiation of CD63^+^SPP1^+^ scar-associated macrophages and liver fibrosis. (A) Representative picro Sirius red staining of olive, CCl_4,_ ALK5-, anti-IL-17A-and anti-GM-CSF-treated livers. (B) Quantification of hydroxyproline liver content. (C) UMAP FACS plot of CD45+ cells. (D) UMAP density plots of CD45+ cells from olive, CCl_4,_ ALK5-, anti-IL-17A-and anti-GM-CSF-treated livers. (E) Stacked bar chart of CD45^+^ cells population in CC_l4_ injury following treatments. (F) Representative FACS plot of CD9 and Trem2 on Ly6C^lo^F4/80^+^ hepatic macrophages. (G) Histogram of CD63 and GPNMB (H) on CD9^+^Trem2^+^ macrophages from olive (grey), CCl_4_ (dark red), ALK5-(light salmon), anti-IL-17A-(salmon) and anti-GM-CSF-treated (light red) livers. (I) Pie-chart representing the frequency of monocytes, Ly6C^hi^ macrophages, CD9^neg^Trem2^neg^ macrophages, CD9^+^Trem2^neg^ macrophages, CD63^lo^ scar-associated macrophages and CD63^hi^ macrophages per treatment group. (J) Quantification of CD63 and GPNMB (K) MFI from olive (grey), CCl_4_ (dark red), ALK5-(light salmon), anti-IL-17A-(salmon) and anti-GM-CSF-treated (light red) livers. (L) Frequencies of CD9^+^Trem2^+^CD63^lo^ and CD9^+^Trem2^+^CD63^hi^ (M) scar-associated macrophages out of CD11b^+^ cells. (N) Frequencies of CD9^+^Trem2^+^CD63^lo^ and CD9^+^Trem2^+^CD63^hi^ (O) scar-associated macrophages out of CD45^+^ cells.

#### Therapeutic TGF-β blockade reduced Fab5 macrophages and fibrosis in lung and liver

We next investigated whether therapeutic TGF-β inhibition with both a small molecule (ALK5i) and a pan-TGF-β neutralizing antibody (1D11) would reduce scar-associated macrophage differentiation in liver and lung fibrosis models. Inhibition of TGF-β led to significant reduction of liver and lung fibrosis (Figure 7A-F). RNA Velocity analysis suggested that 1D11 treatment reduced differentiation of monocytes to Spp1^+^ scar-associated macrophages (Fig. 7G-H). We validated the decrease in scar-associated macrophages by flow cytometry (Fig. 7I). There was a decrease in CD9^+^TREM2^+^CD63^lo^ macrophages in both models of liver fibrosis, but no change in the lung (Fig. 7J, L, N). Importantly, there was a significant decrease in CD9^+^TREM2^+^CD63^hi^ macrophages in the relevant organ for each model, identifying this population as the best correlate with the extent of fibrosis (Fig. 7K, M, O). Using cycIF, we demonstrated that while there was no decrease in SPP1^+^ single-positive cells, there was a highly significant reduction in SPP1^+^GPNMB^+^FABP5^+^ triple-positive cell frequency and area with TGF-β inhibition (Fig. 7P-Q). This result reinforced the earlier finding that a combination of markers is critical for accurate identification of scar-associated macrophages across organs and species, and demonstrated that TGF-β is important for SAM differentiation in both liver and lung fibrosis.

**Figure 7.**
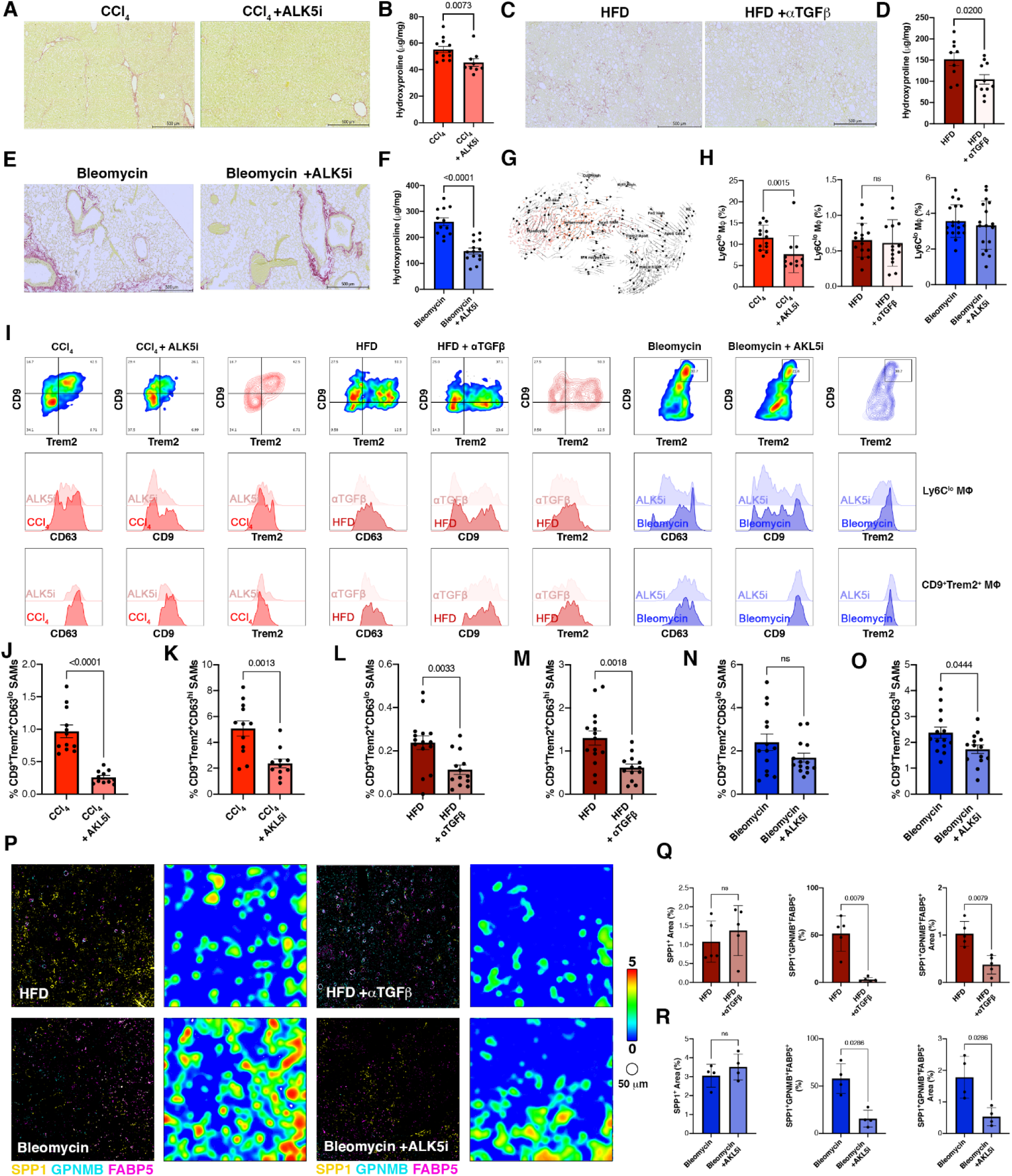
Therapeutic TGF-β blockade inhibits differentiation of SPP1^+^ scar-associated macrophages and fibrosis in murine models of liver and lung disease. (A) Representative picro Sirius red staining of CCl_4_ and ALK5i-treated livers. (B) Quantification of hydroxyproline content in CCl_4_ injury. (C) Representative picro Sirius red staining of HFD and anti-TGF-β (1D11) treated livers. (D) Quantification of hydroxyproline content in HFD. (E) Representative picro Sirius red staining of bleomycin and ALK5i-treated lungs. (F) Quantification of hydroxyproline content in bleomycin lung injury. (G) RNA velocity field projected onto a UMAP plot of myeloid cells from HFD mice treated with a TGF-β blocking antibody. (H) Frequency of Ly6C^lo^CD68^+^F4/80^+^ macrophages in CCl_4_ (red), CCl_4_ + ALK5i (salmon), HFD (dark red), HFD + anti-TGF-β (bordeau), bleomycin (blue) and bleomycin + ALK5i (light blue) by FACS. (I) Representative FACS plots of CD9 and Trem2 on Ly6C^lo^F4/80^+^ macrophages, and histograms of CD63, CD9 and Trem2 on Ly6C^lo^F4/80^+^ macrophages and CD9^+^Trem2^+^ scar-associated macrophages. (J-O) Frequencies of CD9^+^Trem2^+^CD63^lo^ (J, L and N) and CD9^+^Trem2^+^CD63^hi^ (K, M and O) scar-associated macrophages in CCl_4_ (red) and CCl_4_ + ALK5i (salmon) in J and K, HFD (dark red) and HFD + anti-TGF-β (bordeau) in L and M, and bleomycin (blue) and bleomycin + ALK5i (light blue) in N and O. (P) CycIF staining of SPP1, GPNMB and FAPB5 from HFD, HFD + anti-TGF-β bleomycin and bleomycin + ALK5i. Representative tissue heatmaps (scale, blue = 0 to red = 5 cells/50 μm) of SPP1^+^ cells. (Q) Total SPP1-positive area (left panel), frequency of SPP1^+^GPNMB^+^FABP5^+^ scar-associated macrophages (middle panel) and area of SPP1^+^GPNMB^+^FABP5^+^ scar-associated macrophages (right panel) in HFD + anti-TGF-β. (R) Total SPP1^+^ area (left panel), frequency of SPP1^+^GPNMB^+^FABP5^+^ scar-associated macrophages (middle panel) and area of SPP1^+^ GPNMB^+^FABP5^+^ scar-associated macrophages (right panel) in bleomycin + ALK5i.

## Discussion

Contributions of macrophages to promoting and resolving fibrosis have been recognized for many years (*10*). Recently, elegant work demonstrated that CD9^+^TREM2^+^ macrophages expand in fibrotic organs (*11, 13, 19*). Here, leveraging publicly available and newly generated single-cell RNA-Seq datasets, we constructed a multi-species, multi-tissue cellular atlas to better understand the complex architecture of fibrotic niches and how this inflammatory environment drives the scar-association phenotype of macrophages. Using this atlas, we identified a CD9^+^TREM2^+^ scar-associated macrophage phenotype that is expanded in human liver and lung fibrosis. We show that, although TREM2 and CD9 have previously been described as a phenotype of interest, the joint expression of these markers is observed in multiple myeloid populations in liver and lung fibrosis (*11–13, 19*). CD9 and TREM2 are also expressed by homeostatic adipose-resident and alveolar macrophages, further supporting the necessity of using a larger set of well-defined markers to specifically identify scar-associated macrophages (*18*). We used our scRNAseq atlases to determine that joint expression of *Spp1*, *Gpnmb*, *Fabp5* and/or *CD63* with *Trem2* and *CD9* was necessary to specifically identify the expanded, scar-associated macrophage population in fibrotic livers and lungs across species, now abbreviated to Fab5, for their five-marker positivity. By employing flow cytometry and cycIF we orthogonally validated this marker combination across species and tissues. Critically, our analyses employing a defined set of markers can now be used to accurately identify this putatively pathogenic macrophage population.

Given the heterogeneity of macrophages, it was unsurprising that the combination of several markers was required to accurately identify expanded scar-associated macrophages displaying reproducible pro-fibrotic activity (*14*). We predict that this level of precision will become the standard in defining additional macrophage populations with unique functional activities. Our need to use several markers ties into another challenge facing macrophage biologists: clear and shared nomenclature and definitions. Scar-associated macrophages have shared marker expression with cells previously dubbed “lipid-associated macrophages” and “NASH-associated macrophages” among others (*11, 13, 21*). As we and others have demonstrated scar-associated macrophages occur regardless of lipid accumulation, which suggests that “lipid-associated macrophages” is too narrow a term. Similarly, because these macrophages were observed in multiple tissues irrespective of NASH status, we think that “NASH-associated macrophages” is too imprecise a term. Instead, we agree with the previous proposal of “scar-associated macrophages” as the overarching name for these cells; because it avoids tissue- and disease-specific connotations, with “lipid-associated macrophages” and “NASH-associated macrophages” as subsets of “scar-associated macrophages” (*10, 13, 21*). Based on observations across tissues and species, we propose that the definition of a “scar-associated macrophage” should include: 1) expansion in a fibrotic tissue; 2) physical proximity to excessive extracellular matrix; and 3) expression of at least five markers among Trem2, CD9, Spp1, Gpnmb, Fabp5 and CD63. Ideally, direct formal proof of a pro-fibrotic role for scar-associated macrophages should be included in the definition, if functionally validated.

Having generated tools to specifically identify scar-associated macrophages we next investigated their origins. Our RNA Velocity analyses suggested that scar-associated macrophages in liver and lung fibrosis are derived from infiltrating monocytes, which supports data from genetic fate-tracing and computational approaches (*11, 12, 20, 21*). We observed mixed inflammatory signals in our pre-clinical models of fibrosis, as in human diseases. However, scars were enriched in IL-17A, GM-CSF, and TGF-β-activating MMP9 and CD29, leading us to hypothesize that type 3 cytokines drive scar-associated macrophage differentiation (*8, 9, 46*). Interestingly, GM-CSF-derived macrophages express higher levels of CD9, TREM2, CD63 and SPP1 when compared to M-CSF-derived macrophages (Figure 5). Furthermore, GM-CSF is a known inducer of SPP1 in myeloid cells (*47*). Type 1 cytokines (IFNγ) and type 2 cytokines both inhibited expression of SPP1 and scar-associated markers. In contrast, the type 3 stimuli TGF-β and IL-17A supported the differentiation of GM-CSF-derived macrophages to a scar-associated like phenotype with elevated expression of SPP1, CD63, GPNMB, CD9, TREM2. SPP1 and FABP5, two of the top genes expressed by scar-associated macrophages, are linked to the regulation of type 3 responses through the induction of Th17 cells (*48, 49*). Furthermore, we demonstrated that therapeutic blockade of either TGF-β, IL-17A or GM-CSF in CCl_4_ led to a significant reduction of fibrosis and differentiation of CD63^+^SPP1^+^ scar-associated macrophages, confirming our *in vitro* observation (Fig. 6). Taken together, these data suggest that scar-associated macrophages are part of a cycle promoting pro-fibrotic type 3 responses, known to regulate fibrosis progression in multiple organs. Finally, *in vivo* inhibition of TGF-β with small or large molecules led to a decrease in fibrosis and as previously reported rebound type 2 inflammation with elevated levels of IL-5 and IL-13 (*8, 50*). Consistent with our *in vitro* data we observed a significant decrease of CD63^+^SPP1^+^ scar-associated macrophages (Fig. 6), which correlated with decreased fibrosis, TGF-β signaling and IL-17A. While our data support that type 3 inflammation and TGF-β are required for the induction of CD63^+^SPP1^+^ scar-associated macrophages, further studies are needed to characterize the phenotype and function of scar-associated macrophages in type 2 cytokine-driven fibrosis.

Multiple lines of evidence support a pro-fibrotic role for these scar-associated macrophages. We showed that scar-associated-like macrophages significantly enhance deposition of collagen I by HSCs *in vitro*. Furthermore, Ramachandran et al. demonstrated that conditioned media from scar-associated macrophages induces expression of *Col1a1* and *Col1a3* in HSCs (*11*). Additionally, scar-associated macrophages express a variety of genes associated with fibrosis progression. Scar-associated macrophages express Cxcl16, which contribute to the pro-inflammatory/fibrotic Cxcr6 axis (*51, 52*). Dudek et al. recently demonstrated that PD1^+^CXCR6^+^ cytotoxic T cells contribute to pathogenesis of NASH (*53*). Interestingly, we found a similar phenotype in our studies and proposed that scar-associated macrophages orchestrate this phenotype. GPNMB is a glycoprotein that contributes to the degradation and recycling of collagen IV, a normal constituent of liver ECM. GPNMB was recently described as genetically associated with fibrosis progression (*54, 55*). DBA2j mice, which lack expression of GPNMB, have delayed fibrosis, lower HSCs activation and lower mRNA levels of *Col1a1* and *Tgfb1* in acute liver injury (*55*). Furthermore, scar-associated macrophages also express cathepsin B, a protease involved in the lysosomal degradation of collagen IV (*56*). We confirmed these observations by demonstrating increased MMPs activity and degradation of normal ECM substrates by *in vitro* differentiated scar-associated macrophages (Fig. 5O-T). This is further supported by the lack of Cathepsin K and L, MMP12 and MMP13, which are involved in the degradation of fibrillar type I collagen (*10, 57*). Located at the edge of the scar, the expression of GPNMB and cathepsin B by Fab5 macrophages suggests that they contribute to the turnover of normal ECM allowing deposition of pathogenic ECM by activated mesenchymal cells. In addition, CD63 contributes to integrin β-1/3 mediated activation of TGF-β and enhances SMAD3 signaling through activation of the β-catenin pathway (*58*). Furthermore, SPP1 and FABP5 positively modulate Th17 responses creating a feed-forward loop which increases levels of IL-17A, GM-CSF and TGF-β in the scar (*48, 49*). These three cytokines amplify the fibrogenic response through the differentiation of monocytes to Fab5 macrophages and activation of myofibroblasts (Fig. 5-7). While the circumstantial evidence is strong, definitive proof remains elusive and the molecular mechanisms require further study.

In conclusion, here we report an unbiased approach to identify and validate markers of a new “core” fibrotic SAM phenotype that is enriched in lung and liver fibrosis across species. We expanded upon previous identification of SAMs in multiple organs and provide a unifying set of SAM markers that work across species, organs and techniques. These markers led to the identification of signaling molecules that help drive SAM differentiation, and to potential mechanisms by which SAMs promote fibrosis. This process also provides a workflow for identification of cell populations associated with human diseases using scRNAseq data, and to then validate the existence of these populations in animal models for the prioritization of therapeutic strategies and targets for future validation. In this publication, we have used this approach to identify a well-defined population of SAMs that represent a promising target population for the treatment of human fibrotic disease. A generalized version of this process could be applied to other human diseases to aid in the development of a wide range of therapies.

