## Supplemental Figures 1-9 for "Identification of a Broadly Fibrogenic Macrophage Subset Induced by Type 3 Inflammation in Human and Murine Liver and Lung Fibrosis"

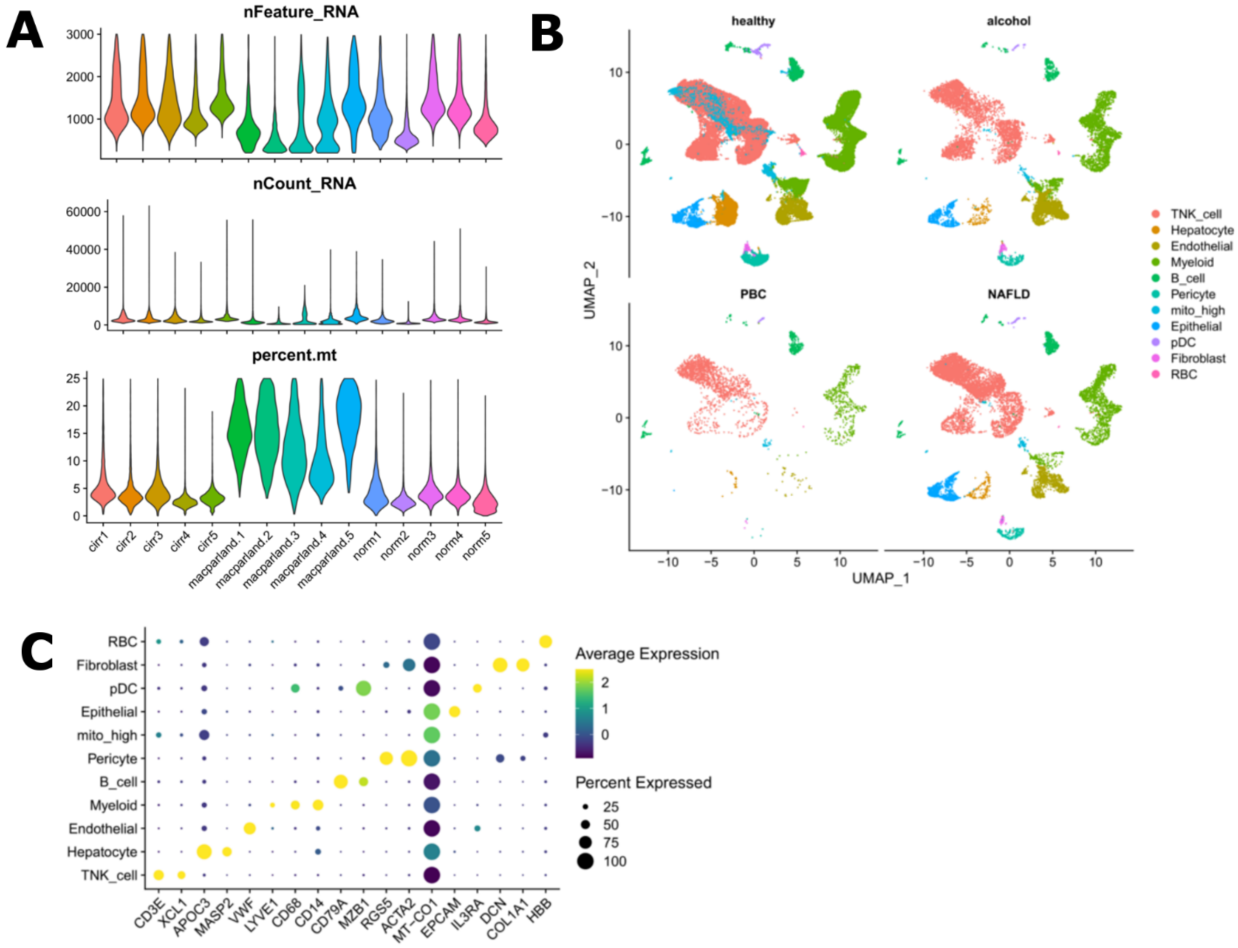


**Supplemental Figure 1. Human liver scRNAseq atlas quality control and annotation metrics.**

(A) scRNAseq quality metrics by patient and study. (B) UMAPs of cell types split by health status. (C) Expression of genes used to annotate clusters with cell type identities. Dot size represents the fraction of cell type (rows) expressing each gene (columns). Hue represents the scaled average expression per gene.

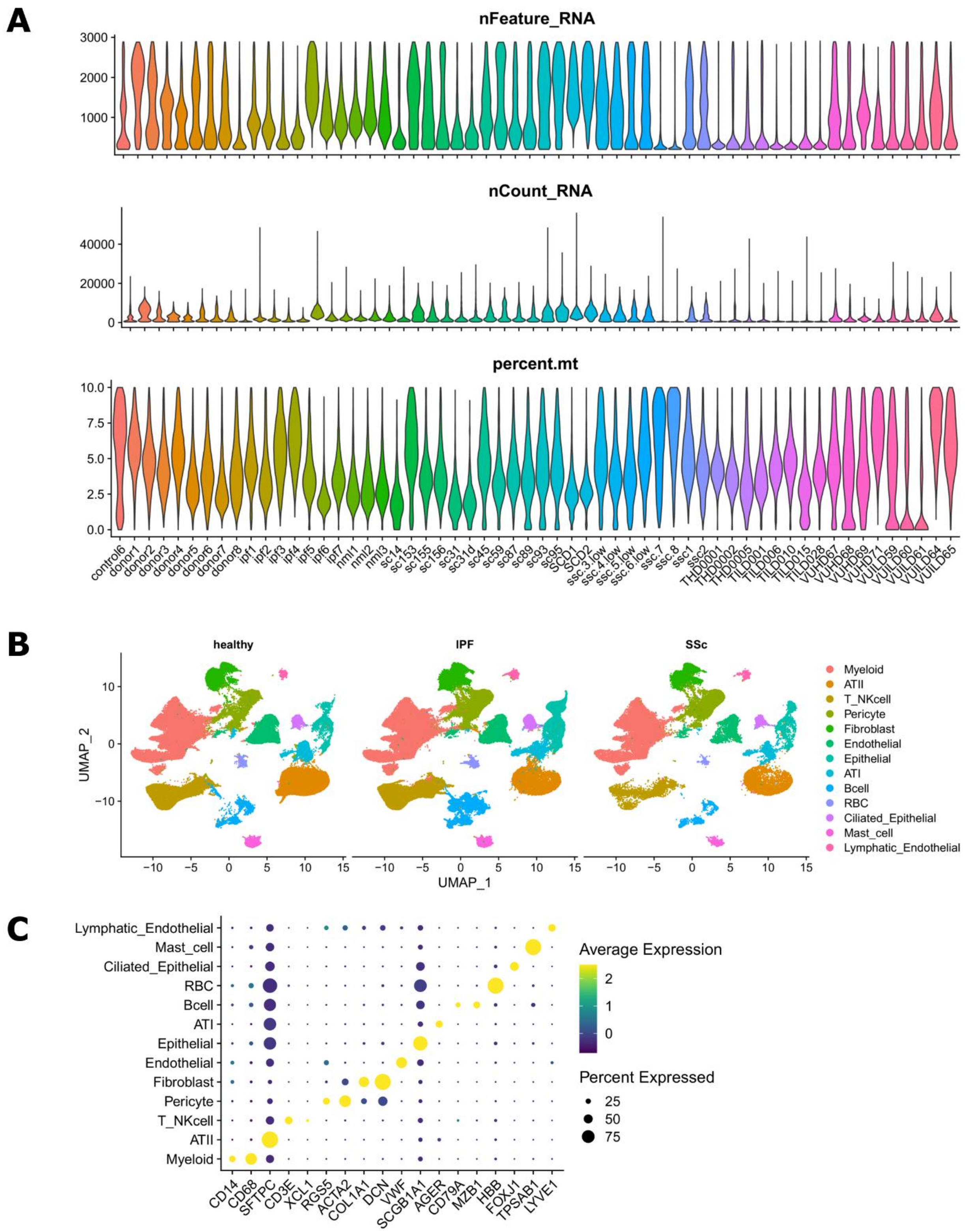


**Supplemental Figure 2. Human lung scRNAseq atlas quality control and annotation metrics.**

(A) scRNAseq quality metrics by patient and study. (B) UMAPs of cell types split by health status. (C) Expression of genes used to annotate clusters with cell type identities. Dot size represents the fraction of cell type (rows) expressing each gene (columns). Hue represents the scaled average expression per gene.

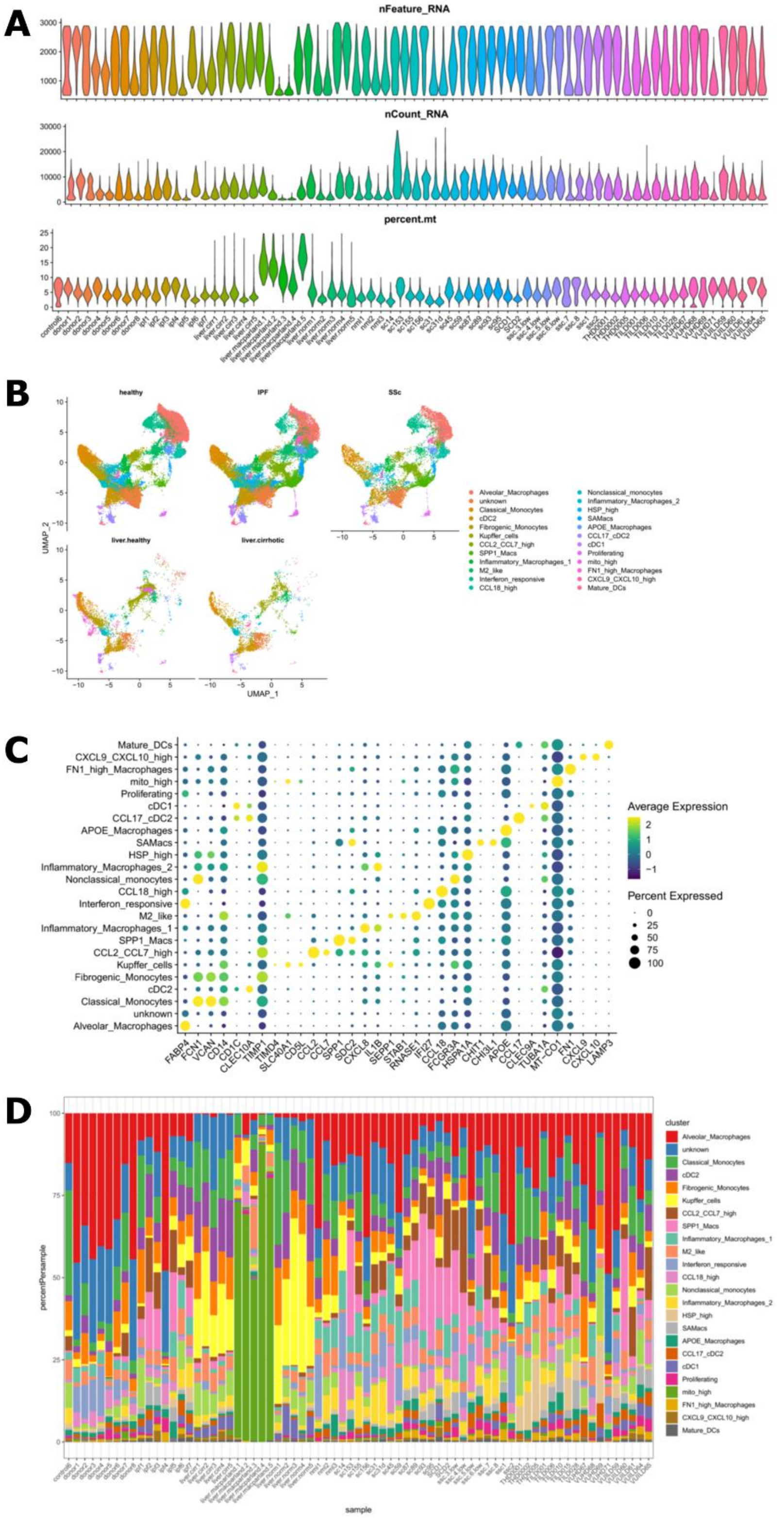


**Supplemental Figure 3. Human myeloid scRNAseq atlas quality control and annotation metrics.**

(A) scRNAseq quality metrics by patient and study. (B) UMAPs of cell types split by health status. (C) Expression of genes used to annotate clusters with myeloid cell identities. Dot size represents the fraction of cell type (rows) expressing each gene (columns). Hue represents the scaled average expression per gene. (D) Proportions of myeloid subsets by patient.

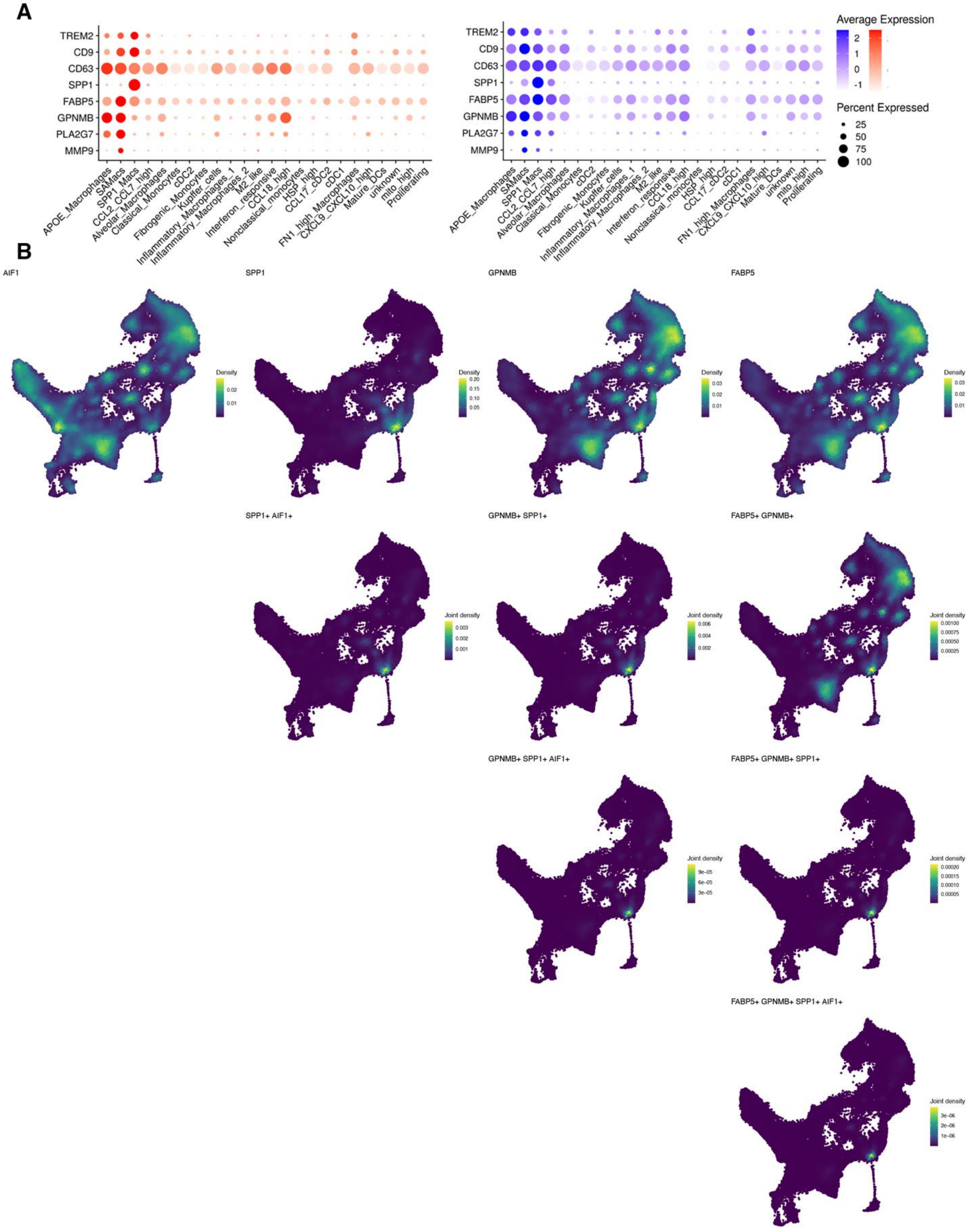


**Supplemental Figure 4. The combination of SPP1, GPNMB and FAPB5 is specific to scar-associated macrophages.**

(A) Dot plot showing expression of scar-associated macrophage genes in myeloid clusters from human livers (red) and lungs (blue). Dot size represents the fraction of cell type (rows) expressing each gene (columns). Hue represents the scaled average expression per gene. (B) Nebulosa plots showing individual (1st row) and joint (2nd-4th rows) densities of AIF1, SPP1, GPNMB and FAPB5.

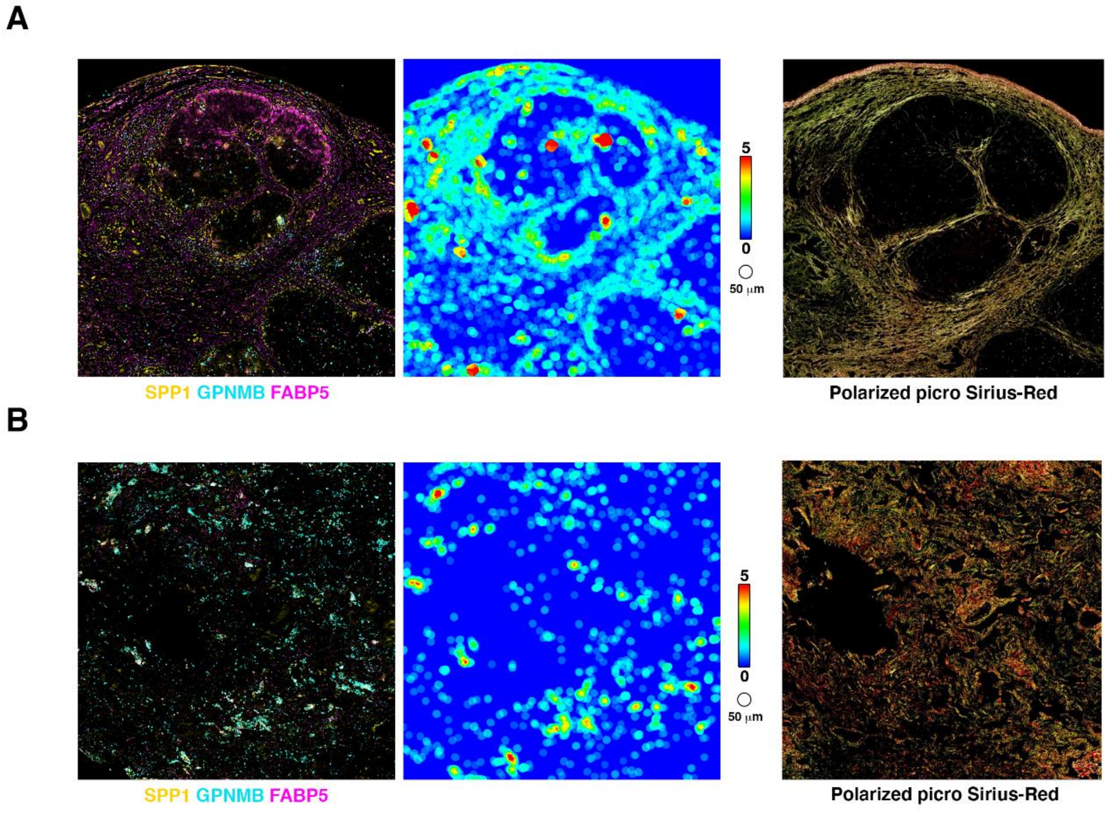


**Supplemental Figure 5. SPP1$^+$ scar-associated macrophages are located at the edges of the scar**

(A) Immunofluorescent staining (left) of SPP1 (yellow), GPNMB (cyan) and FAPB5 (magenta), tissue heatmap (middle) of SPP1$^+$ cells and polarized picro Sirius-red images (right) of a NASH F4 liver. (B) Immunofluorescent staining (left) of SPP1 (yellow), GPNMB (cyan) and FAPB5 (magenta), tissue heatmap (middle) of SPP1$^+$ cells and polarized picro Sirius-red images (right) of a fibrotic IPF lung.

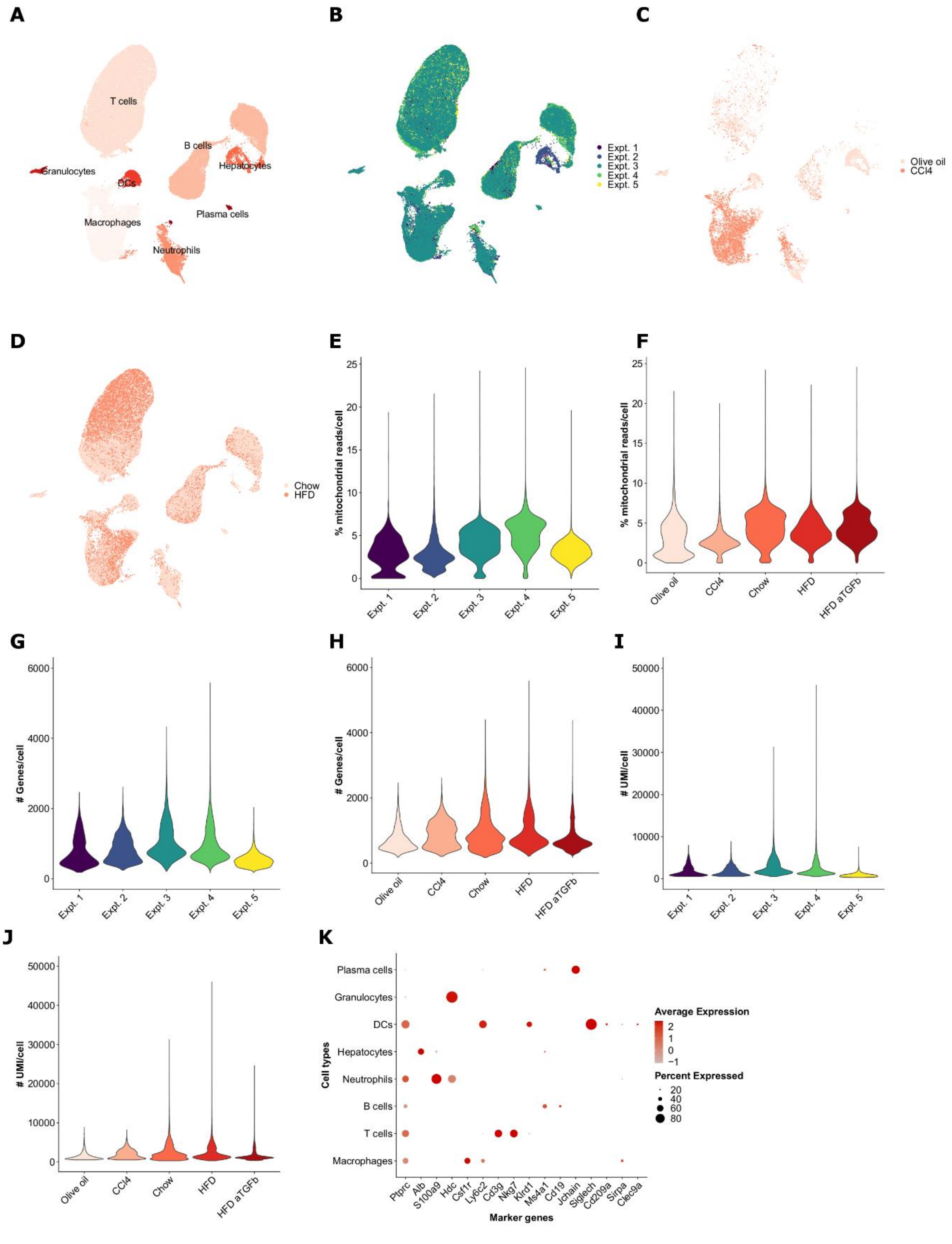


**Supplemental Figure 6. Murine liver scRNAseq quality control and annotation metrics.**

(A) Annotated UMAP of all sequenced intrahepatic cells colored by cell type after filtering low quality cells and multiplets. (B) Same UMAP as in (A) colored by experiment. (C) Subset of cells from panels (A) & (B) from mice treated with olive oil or $CCl_4$ colored by treatment. (D) Subset of cells from panels (A) & (B) from mice fed normal chow or the GAN diet (HFD) colored by diet. (E-J) Single-cell quality metrics by experiment (E, G, I) or treatment/diet (F, H, J). (K) Expression of genes used to annotate clusters with cell type identities. Dot size represents the fraction of cell type (rows) expressing each gene (columns). Hue represents the scaled average expression per gene. For ease of interpretation dot.min=0.15 so that dots are shown only for clusters where at least 15% of cells express the indicated gene.

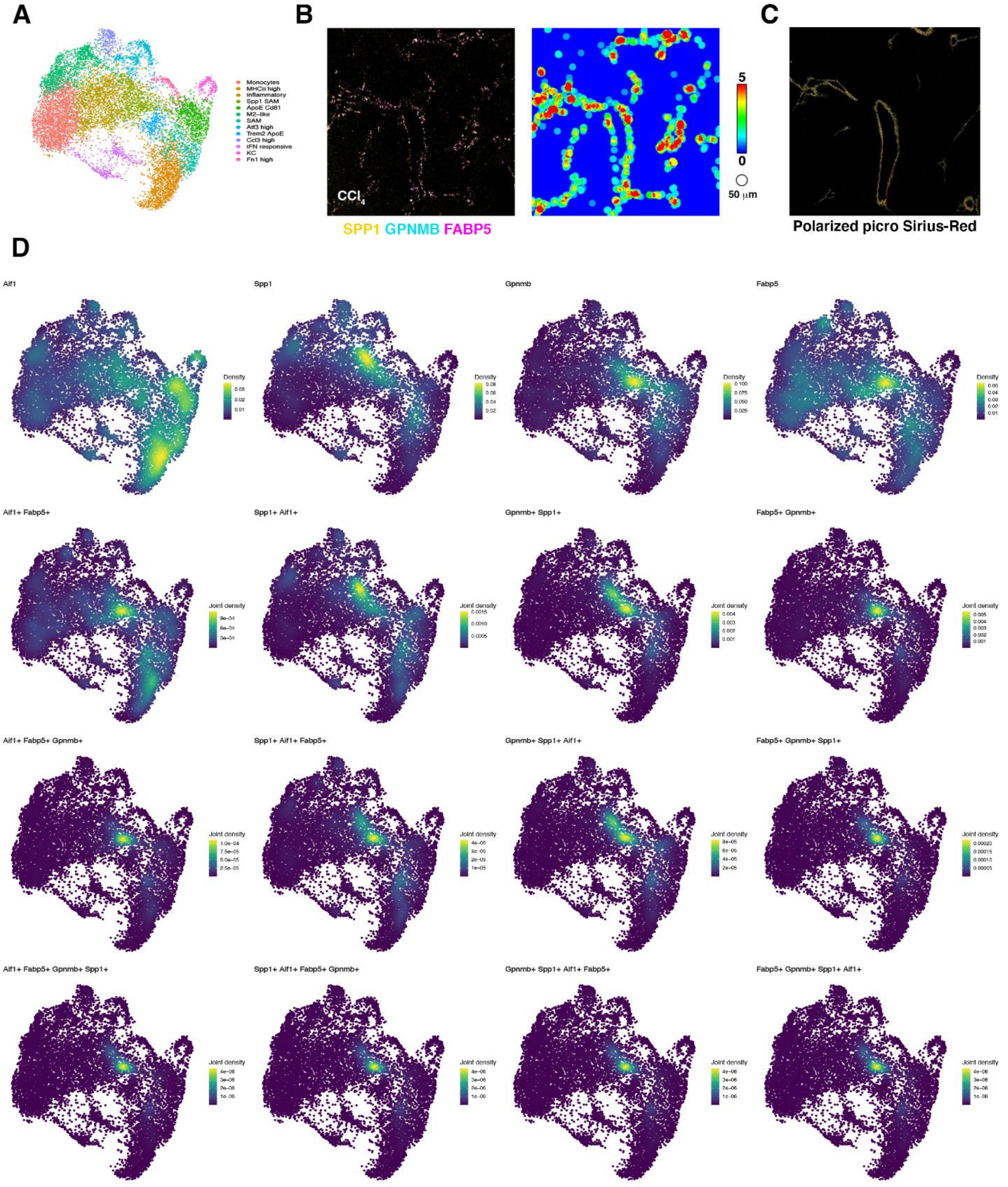


**Supplemental Figure 7. Murine SPP1$^{+}$ scar-associated macrophages are also identified by the co-expression of SPP1, GPNMB and FABP5 and are located at the edges of the scar.**

(A) scRNAseq UMAP of myeloid cells from healthy, $CCl_4$ and HFD mouse livers with cell type annotations. (B) Immunofluorescent staining (left) of SPP1 (yellow), GPNMB (cyan) and FAPB5 (magenta), tissue heatmap (right) of $SPP1^+$ cells. (C) Polarized picro Sirius-red image (right) of a chronic $CCl_4$ liver section (immediately adjacent section to that in (B)). (D) Nebulosa plots showing individual (1st row) and joint (2nd-4th row) densities of AIF1, SPP1, GPNMB and FAPB5.

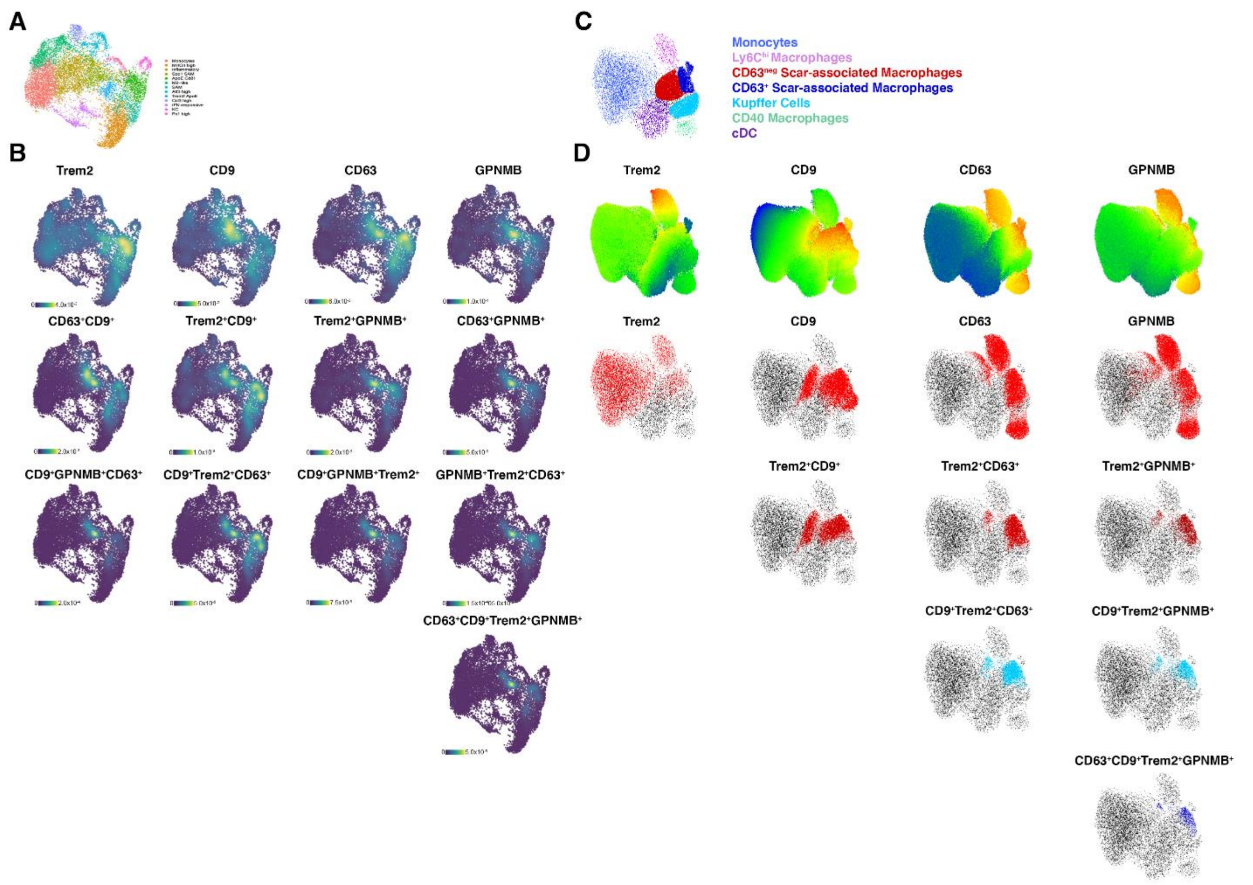


**Supplemental Figure 8. Combining TREM2, CD9, CD63 and GPNMB enhanced the precise identification of SPP1$^{+}$ scar-associated macrophages by scRNAseq and flow cytometry.**

(A) scRNAseq UMAP of myeloid cells from healthy, $CCl_4$ and HFD mouse livers with clusters annotated by subset. (B) Nebulosa plots showing individual (1$^{st}$ row) and joint (2$^{nd}$-4$^{th}$ rows) densities of *Trem2*, *Cd9*, *Cd63* and *Gpnmb*. (C) FACS UMAP of myeloid cells from healthy, $CCl_4$ and HFD mouse livers with clusters annotated by subset. (D) Top row: Heatmap of TREM2, CD9, CD63 or GPNMB protein expression (blue represents low and red represents high expression) on the UMAP from (C). Remaining rows: single (2$^{nd}$ row) or joint (3$^{rd}$-5$^{th}$ rows) distributions of myeloid cells positive for TREM2, CD9, CD63 and/or GPNMB.

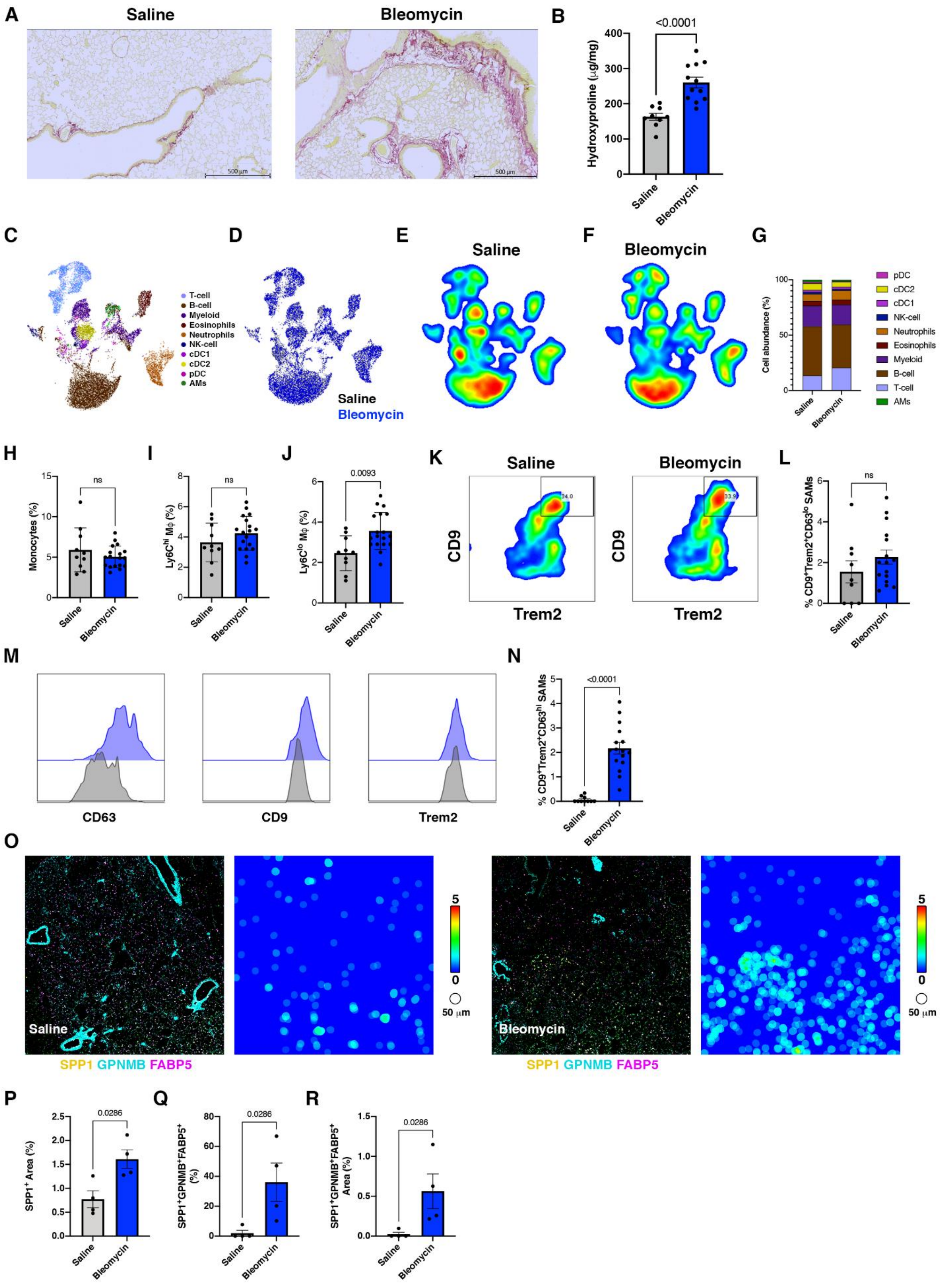
A
Saline
Bleomycin
500 µm
500 µm
B
<0.0001
Hydroxyproline (µg/mg)
400
300
200
100
0
Saline
Bleomycin
C
T-cell
B-cell
Myeloid
Eosinophils
Neutrophils
NK-cell
cDC1
cDC2
pDC
AMs
D
Saline
Bleomycin
E
Saline
F
Bleomycin
G
Cell abundance (%)
100
50
0
Saline
Bleomycin
pDC
cDC2
cDC1
NK-cell
Neutrophils
Eosinophils
Myeloid
B-cell
T-cell
AMs
H
ns
Monocytes (%)
15
10
5
0
Saline
Bleomycin
I
ns
Ly6C^hi Mφ (%)
8
6
4
2
0
Saline
Bleomycin
J
0.0093
Ly6C^lo Mφ (%)
6
4
2
0
Saline
Bleomycin
K
Saline
Bleomycin
CD9
Trem2
34.0
33.9
CD9
Trem2
L
ns
% CD9+Trem2+CD63lo SAMs
6
4
2
0
Saline
Bleomycin
M
CD63
CD9
Trem2
N
<0.0001
% CD9+Trem2+CD63hi SAMs
5
4
3
2
1
0
Saline
Bleomycin
O
Saline
SPP1 GPNMB FABP5
5
0
50 µm
Bleomycin
SPP1 GPNMB FABP5
5
0
50 µm
P
0.0286
SPP1+ Area (%)
2.5
2.0
1.5
1.0
0.5
0.0
Saline
Bleomycin
Q
0.0286
SPP1+GPNMB+FABP5+ (%)
80
60
40
20
0
Saline
Bleomycin
R
0.0286
SPP1+GPNMB+FABP5+ Area (%)
1.5
1.0
0.5
0.0
Saline
Bleomycin

**Supplemental Figure 9. The frequency of SPP1$^+$ scar-associated macrophages is increased in lung fibrosis**

(A) Representative picro Sirius-red staining of saline and bleomycin treated lungs. (B) Hydroxyproline quantification from saline and bleomycin treated lungs. (C) UMAP FACS plots of CD45$^+$ cells from bleomycin injury. (D) UMAP overlay of saline (grey) and bleomycin (blue). UMAP FACS density plots of CD45$^+$ cells in saline (E) and bleomycin (F). (G) Stacked bar chart of CD45$^+$ cells population in bleomycin lung injury. Frequency of Ly6C$^{hi}$ monocytes (H), Ly6C$^{hi}$CD68$^+$F4/80$^+$ macrophages (I) and Ly6C$^{lo}$CD68$^+$F4/80$^+$ macrophages (J). (K) Representative FACS plot of CD9 and Trem2 on Ly6C$^{lo}$F4/80$^+$ pulmonary macrophages of saline (left) and bleomycin (right). (L) Frequency of CD9$^+$Trem2$^+$CD63$^{lo}$ scar-associated macrophages. (M) Histograms of CD63, CD9 and Trem2 expression by FACS on CD9$^+$Trem2$^+$ scar-associated macrophages isolated from saline (grey) and bleomycin (blue) lungs. (N) Frequency of CD9$^+$Trem2$^+$CD63$^{hi}$ scar-associated macrophages. (O) CycIF staining of SPP1, GPNMB and FAPB5 from saline and bleomycin lungs (left) and representative tissue heatmaps (right, scale: blue = 0 to red = 5 cells/50 $\mu$m) of SPP1$^+$ cells. (P) Total SPP1$^+$ area by CycIF in lung sections. (Q) Frequency of SPP1$^+$GPNMB$^+$FABP5$^+$ scar-associated macrophages by CycIF in lung sections. (R) Area of SPP1$^+$GPNMB$^+$FABP5$^+$ scar-associated macrophages by CycIF in lung sections. Statistics Mann-Whitney u test, p-values displayed on plots.
