## Supplemental Table 1 for "Identification of a Broadly Fibrogenic Macrophage Subset Induced by Type 3 Inflammation in Human and Murine Liver and Lung Fibrosis"

| Patient ID # | Age | Sex | Race | Tissue | Condition | FibroTest/FibroSure Score | FEV1 | FVC | TLC | TLCO | KCO | Pack yrs |
|---|---|---|---|---|---|---|---|---|---|---|---|---|
| 1 | 61 | F | White | Liver | NASH | 2 | N/A | N/A | N/A | N/A | N/A | N/A |
| 2 | 51 | M | White | Liver | NASH | 3 | N/A | N/A | N/A | N/A | N/A | N/A |
| 3 | 69 | F | White | Liver | NASH | 4 | N/A | N/A | N/A | N/A | N/A | N/A |
| 4 | 48 | M | White | Liver | NASH | 1 | N/A | N/A | N/A | N/A | N/A | N/A |
| 5 | 56 | F | White | Liver | NASH | 2 | N/A | N/A | N/A | N/A | N/A | N/A |
| 6 | 81 | F | White | Liver | NASH | 1 | N/A | N/A | N/A | N/A | N/A | N/A |
| 7 | 74 | M | White | Liver | NASH | 0 | N/A | N/A | N/A | N/A | N/A | N/A |
| 8 | 34 | F | White | Liver | NASH | 3 | N/A | N/A | N/A | N/A | N/A | N/A |
| 9 | 34 | F | White | Liver | NASH | 3 | N/A | N/A | N/A | N/A | N/A | N/A |
| 10 | 56 | F | American Indian | Liver | NASH | 3 | N/A | N/A | N/A | N/A | N/A | N/A |
| 11 | 55 | M | White | Liver | NASH | 3 | N/A | N/A | N/A | N/A | N/A | N/A |
| 12 | 55 | M | White | Liver | NASH | 3 | N/A | N/A | N/A | N/A | N/A | N/A |
| 13 | 58 | F | White | Liver | NASH | 3 | N/A | N/A | N/A | N/A | N/A | N/A |
| 14 | 56 | F | White | Liver | NASH | 1 | N/A | N/A | N/A | N/A | N/A | N/A |
| 15 | 56 | F | White | Liver | NASH | 1 | N/A | N/A | N/A | N/A | N/A | N/A |
| 16 | 76 | M | White | Liver | NASH | 1 | N/A | N/A | N/A | N/A | N/A | N/A |
| 17 | 76 | M | White | Liver | NASH | 1 | N/A | N/A | N/A | N/A | N/A | N/A |
| 18 | 72 | M | White | Liver | NASH | 1 | N/A | N/A | N/A | N/A | N/A | N/A |
| 19 | 64 | M | White | Liver | NASH | 1 | N/A | N/A | N/A | N/A | N/A | N/A |
| 20 | 66 | M | White | Liver | NASH | 3 | N/A | N/A | N/A | N/A | N/A | N/A |
| 21 | 68 | M | White | Liver | NASH | 3 | N/A | N/A | N/A | N/A | N/A | N/A |
| 22 | 28 | M | N/A | Lung | Healthy | N/A | N/A | N/A | N/A | N/A | N/A | Smoker |
| 23 | 42 | M | N/A | Lung | Healthy | N/A | N/A | N/A | N/A | N/A | N/A | Non-smoker |
| 24 | 40 | M | N/A | Lung | Healthy | N/A | N/A | N/A | N/A | N/A | N/A | Ex-smoker - stopped Nov 2017 |
| 25 | 52 | F | N/A | Lung | Healthy | N/A | N/A | N/A | N/A | N/A | N/A | Smoker, Cause of Death intracranial thrombosis |
| 26 | 52 | F | N/A | Lung | Healthy | N/A | N/A | N/A | N/A | N/A | N/A | Non-smoker |
| 27 | N/A | N/A | N/A | Lung | Healthy | N/A | N/A | N/A | N/A | N/A | N/A | N/A |
| 28 | N/A | N/A | N/A | Lung | Healthy | N/A | N/A | N/A | N/A | N/A | N/A | N/A |
| 29 | N/A | N/A | N/A | Lung | Healthy | N/A | N/A | N/A | N/A | N/A | N/A | N/A |
| 30 | 58 | M | N/A | Lung | IPF | N/A | 2.25 (70%) | 2.60 (63%) | 3.97 (61%) | 5.91 (65%) | 1.68 (116%) | Non smoker |
| 31 | 65 | M | N/A | Lung | IPF | N/A | 2.18 (63%) | 2.60 (58%) | 4.42 (59%) | 2.79 (28%) | 0.66 (50%) | Stopped smoking 2013 only smoked 1-2 cigarettes a day |
| 32 | 62 | F | N/A | Lung | IPF | N/A | 0.80 (30%) | 1.25 (40%) | 2.79 (51%) | 2.76 (33%) | 1.37 (90%) | Non Smoker |
| 33 | 59 | M | N/A | Lung | IPF | N/A | 2.02 (57%) | 2.18 (48%) | 4.19 (57%) | 5.00 (49%) | 1.42 (102%) | Smoking 20 pack yrs, stopped 12 yr ago. |
| 34 | 62 | M | N/A | Lung | IPF | N/A | 1.16 (37%) | 1.57 (40%) | 2.49 (37%) | 1.06 (12%) | 0.55 (41%) | Ex- smoker stopped 2014 34 pack years. |
| 35 | 59 | M | N/A | Lung | IPF | N/A | 2.35 (54%) | 2.76 (54%) | 3.15 (39%) | 3.44 (30%) | 0.98 (71%) | Ex-smoker 30 pack years |
| 36 | 63 | F | N/A | Lung | IPF | N/A | 1.55 (67%) | 1.65 (58%) | Not recorded | 2.75 (36%) | 1.17 (77%) | Never smoked |
| 37 | 53 | M | N/A | Lung | IPF | N/A | 2.96 (76%) | 4.27 (88%) | 7.18 (94%) | 3.54 (32%) | 0.75 (52%) | Ex-Smoker stopped 2013, 30 pack year |
| 38 | 61 | M | N/A | Lung | IPF | N/A | 1.52 (50%) | 1.7 (44%) | 2.51 (39%) | 2.42 (27%) | 1.02 (75%) | Ex smoker stopped 2005 |
| 39 | 66 | M | N/A | Lung | IPF | N/A | 2.47 (81%) | 2.86 (73%) | 4.16 (62%) | 2.88 (33%) | 0.81 (61%) | Non-smoker |
| 40 | 62 | M | N/A | Lung | IPF | N/A | 2.21(69%) | 2.72(66%) | 2.94(32%) | 2.94(32%) | 0.80(59%) | Ex smoker stopped 2011 40-50 pack years |
| 41 | 56 | M | N/A | Lung | IPF | N/A | 1.53(44%) | 2.55(58%) | 4.37(61%) | 3.48(53%) | 0.95(68%) | Ex-smoker stopped 1998, 20 pack years |
| 42 | 51 | F | N/A | Lung | IPF | N/A | 1.67(63%) | 1.85(55%) | 2.45(48%) | 3.13(38%) | 1.36(83%) | Ex-smoker stopped 2012 30 pack yr |
| 43 | 64 | M | N/A | Lung | IPF | N/A | 2.29(64%) | 3.02(66%) | 4.66(61%) | 2.87(28%) | 0.83(62%) | Ex-smoker stopped 2015 40 pack yr |
| 44 | 59 | F | N/A | Lung | IPF | N/A | 2.10(82%) | 2.44(81%) | 5.45(104%) | 1.70(21%) | 0.58(38%) | Ex-smoker stopped Feb 2015 30pk |
| 45 | 64 | M | N/A | Lung | IPF | N/A | 2.32 (72%) | 2.66 (65%) | 4.26 (62%) | 1.75 (19%) | 1.60(65%) | Ex-smoker stopped 2012 30 pack yr |
| 46 | 61 | M | N/A | Lung | IPF | N/A | 2.1 (64%) | 2.32(56%) | 3.32(48%) | 1.76(19%) | 0.58(42%) | Ex-smoker. 15 pack year |
| 47 | 50 | F | N/A | Lung | IPF | N/A | 1.33(46%) | 1.51(45%) | 2.21(41%) | 2.21(25%) | 1.20(75%) | Ex-smoker |

| | | | | | | | | | | | | |
|---|---|---|---|---|---|---|---|---|---|---|---|---|
| 48 | 63 | M | N/A | Lung | IPF | N/A | 2.29 | 2.72 | 4.97 | 7.56 | 1.52 | Unknown if previous smoker |
| 49 | 63 | M | N/A | Lung | IPF | N/A | 2.61 | 3.30 | 5.78 | 7.70 | 1.33 | ex-smoker stoped 2017 |
