## Supplemental Table 2 for "Identification of a Broadly Fibrogenic Macrophage Subset Induced by Type 3 Inflammation in Human and Murine Liver and Lung Fibrosis"

| Antigen | Color | Clone | Company | Dilution | Target Species |
|---|---|---|---|---|---|
| CD3e | BUV395 | 145-2C11 | BD Biosciences | 1:100 | Mouse |
| CD4 | BUV496 | RM4-5 | BD Biosciences | 1:100 | Mouse |
| CD8 | BUV737 | 5H10-1 | BD Biosciences | 1:100 | Mouse |
| TCRgd | BV421 | GL3 | Biolegend | 1:100 | Mouse |
| CD11b | BV605 | M1/70 | Biolegend | 1:100 | Mouse |
| B220 | BV786 | RA3-6B2 | BD Biosciences | 2:100 | Mouse |
| F4/80 | FITC | BM8 | Biolegend | 2:100 | Mouse |
| CD11c | PerCP/Cy5.5 | N418 | Biolegend | 2:100 | Mouse |
| Siglec-F | PE | E50-2440 | BD Biosciences | 2:100 | Mouse |
| Ly6G | PE/Dazzle 594 | 1A8 | Biolegend | 1:200 | Mouse |
| Ly6C | Pe/Cy7 | HK1.4 | Biolegend | 1:100 | Mouse |
| CD68 | APC | FA-11 | Biolegend | 1:100 | Mouse |
| CD45 | A700 | I3/2.3 | Biolegend | 1:100 | Mouse |
| NK1.1 | APC/Fire 750 | S17016D | Biolegend | 2:100 | Mouse |
