## Supplemental Table 3 for "Identification of a Broadly Fibrogenic Macrophage Subset Induced by Type 3 Inflammation in Human and Murine Liver and Lung Fibrosis"

| Antigen | Color | Clone | Company | Dilution | Target Species |
|---|---|---|---|---|---|
| CD3 | BUV395 | 500A2 | BD Biosciences | 1:100 | Mouse |
| B220 | BUV395 | RA3-6B2 | BD Biosciences | 2:100 | Mouse |
| NK1.1 | BUV395 | PK136 | BD Biosciences | 2:100 | Mouse |
| Siglec-F | BUV395 | E50-2440 | BD Biosciences | 1:100 | Mouse |
| CD38 | BUV496 | 90/CD38 | BD Biosciences | 1:50 | Mouse |
| CD11b | BUV737 | M1/70 | BD Biosciences | 2:100 | Mouse |
| CD81 | BV421 | Eat2 | BD Biosciences | 1:100 | Mouse |
| F4/80 | BV605 | BM8 | Biolegend | 2:100 | Mouse |
| CD40 | BV786 | 1C10 | ThermoFisher | 2:100 | Mouse |
| Trem2 | FITC | 2B10C42 | Biolegend | 3:100 | Mouse |
| CD63 | PerCPCy5.5 | NVG-2 | Biolegend | 3:100 | Mouse |
| Ly6G | PE-TxR | HK1.4 | Biolegend | 1:200 | Mouse |
| Ly6C | PE-Cy7 | 1A8 | Biolegend | 1:100 | Mouse |
| GPNMB | APC | TR3MBL1 | ThermoFisher | 3:100 | Mouse |
| CD45 | A700 | 30-F11 | Biolegend | 2:100 | Mouse |
| CD9 | APC-F750 | MZ3 | Biolegend | 3:100 | Mouse |
