## Supplemental Table 4 for "Identification of a Broadly Fibrogenic Macrophage Subset Induced by Type 3 Inflammation in Human and Murine Liver and Lung Fibrosis"

| Antigen | Color | Clone | Company | Dilution | Target Species |
|---|---|---|---|---|---|
| CD45 | AF700 | HI30 | BD Biosciences | 1:200 | Human |
| CD14 | BUV737 | M5E2 | BD Biosciences | 1:200 | Human |
| CD16 | BUV496 | 3G8 | BD Biosciences | 1:200 | Human |
| GPNMB | PE | HOST5DS | ThermoFisher | 1:200 | Human |
| CD9 | PE-Tx | M-L13 | BD Biosciences | 1:200 | Human |
| CD11b | BV786 | D12 | BD Biosciences | 1:200 | Human |
| Trem2 | APC | 237920 | R&D | 2:100 | Human |
| CD63 | PE-Cy7 | H5C6 | Biolegend | 1:200 | Human |
| CD29 | BUV395 | 4-Mar | BD Biosciences | 1:200 | Human |
| CD206 | BV421 | 15-2 | Biolegend | 1:200 | Human |
| CD86 | BV605 | BU63 | Biolegend | 1:200 | Human |
