## Supplemental Table 6 for "Identification of a Broadly Fibrogenic Macrophage Subset Induced by Type 3 Inflammation in Human and Murine Liver and Lung Fibrosis"

**Murine CycIF Antibodies**

| Target | Clone | Catalog # | Company | Dilution |
|---|---|---|---|---|
| Iba1 | pAb | 019-19741 | FujiFilm Wako | 1:50 |
| Spp1 | pAb | AF808 | R&D Systems | 1:50 |
| Fabp5 | pAb | AF1476 | R&D Systems | 1:50 |
| Gpnmb | pAb | NBP1-69389 | Novus | 1:50 |
