## Supplemental Table 7 for "Identification of a Broadly Fibrogenic Macrophage Subset Induced by Type 3 Inflammation in Human and Murine Liver and Lung Fibrosis"

**CycIF Secondary Antibodies**

| Target | Fluor | Catalog # | Company | Dilution |
|---|---|---|---|---|
| goat IgG | AF555 | A-21432 | ThermoFisher | 1:200 |
| rabbit IgG | AF647 | A-31573 | ThermoFisher | 1:200 |
| mouse IgG | AF488 | A-21202 | ThermoFisher | 1:200 |
| rat IgG | AF647 | A48272 | ThermoFisher | 1:200 |
| goat IgG | AF647 | A-21447 | ThermoFisher | 1:200 |
| rabbit IgG | AF555 | A-31572 | ThermoFisher | 1:200 |
